# A memory element imposes epigenetic behavior on intrinsically labile RNAi-induced heterochromatin spread

**DOI:** 10.1101/237479

**Authors:** R.A. Greenstein, Stephen K. Jones, Eric C. Spivey, James R. Rybarski, Ilya J. Finkelstein, Bassem Al-Sady

## Abstract

The heterochromatin spreading reaction is a central contributor to the formation of gene-repressive structures, which are re-established with high fidelity following replication. The high fidelity of this process is not obviously encoded in the primary spreading reaction. To resolve origins of stable inheritance of repression, we probed the intrinsic fidelity of spreading events in fission yeast using a system that quantitatively describes the spreading reaction in live single cells. We show that spreading from RNAi-nucleated elements is stochastic, multimodal, and fluctuates dynamically across time. In contrast, a second form of spreading, nucleated by the *cis*-acting element *REIII*, is deterministic, has high memory capacity and acts as the source of locus fidelity. *REIII* enables fidelity in part by endowing the locus with resistance to perturbations. Together, our results suggest that epigenetic capacity may not be intrinsically encoded in the spreading reaction, but rather requires collaboration with specialized memory elements.

## INTRODUCTION

The formation of gene-repressive heterochromatin domains is critical for genome integrity and for the establishment and maintenance of cell identity. Most heterochromatin formation occurs by a sequence-indifferent spreading reaction that propagates heterochromatic marks, structural proteins, and associated effector proteins outwards from nucleation sites. While this reaction can be treated like the formation of a template-guided polymer (chromatin), it differs from other cellular polymers because the precise extent of its formation has critical heritable consequences for cell identity. For example, in early pluripotent precursors, pre-existing heterochromatin domains spread, sometimes over megabases, to repress specifiers of inappropriate cell types. Importantly, the final extent of spreading from a locus appears to be dependent on the lineage pathway and thus varies across different precursors (Wen et al., 2009; Zhu et al., 2013). Imprecise spreading within a lineage can lead to differentiation defects or disease (Ceol et al., 2011). Similarly, spreading also specifies cell type in yeasts, where the cell type is maintained by repressing the mating cassettes at the mating type loci (Ekwall et al., 1991). Despite the centrality of the spreading reaction in shaping cell identity, its native and intrinsic cellular characteristics, as well as mechanisms for its inter-generational propagation, have remained opaque.

We have some understanding of how cells inherit silencing at nucleation sites, i.e. the DNA-sequence driven component of heterochromatin. Recent results in heterochromatin systems signaled by Histone 3 Lysine 9 and Lysine 27 methylation (H3K9me and H3K27me) indicate that several mechanisms act together to ensure intergenerational inheritance: continuous DNA-mediated recruitment of the histone methylase (Audergon et al., 2015; Laprell et al., 2017; Ragunathan et al., 2015; Wang and Moazed, 2017) low histone turnover (Taneja et al., 2017) as well as the positive “read-write” feedback loop for histone methylases. Additionally, studies suggest that either the histone mark (Gaydos et al., 2014) or the histone methylases (Petruk et al., 2012) can persist trans-generationally.

These insights concerning nucleation sites do not necessarily account for how regions of heterochromatin distal to these sites are maintained at high fidelity. Unlike nucleation, which depends on DNA based enzyme recruitment (Bayne et al., 2010; Verdel et al., 2004), spreading depends on the ability of the system to propagate along the chromosome, independent of the underlying DNA sequence. Such propagation requires the “read-write” positive feedback function of the system (Al-Sady et al., 2013; Margueron et al., 2009; Muller et al., 2016; Noma et al., 2004; Zhang et al., 2008). The reliance on DNA-sequence indifferent spreading for propagating the heterochromatic state takes on special importance in situations where modified nucleosomes are less likely to persist. This is the case in the fission yeast H3K9me-signaled system. Like other model systems, it lacks DNA methylation and, additionally, even antagonizes persistence of the modified state. This is due to the presence of a putative H3K9me histone demethylase, Epe1, which rapidly erases H3K9 methylation, and thus the heterochromatic state (Audergon et al., 2015; Ragunathan et al., 2015). Therefore, the domain must be re-formed by spreading from the original nucleation site every cell cycle (Chen et al., 2008), leaving unresolved the question of how high-fidelity formation of heterochromatin structures is accomplished.

Resolving this question is further complicated by the lack of consensus on the intrinsic character of the spreading reaction in cells. It is not certain whether spreading is deterministic (where spreading is executed to its maximal extent every time nucleation is successful) or stochastic (where only some nucleation events result in a spreading event). In either case, the outcome may or may not yield intrinsic stability, where the heterochromatin state persists through divisions. Both hypotheses have support: For example, early experiments indicated that genetically-disrupted heterochromatin domains are stochastic in their nucleation or spreading behavior in both flies and in fission yeast (Elgin and Reuter, 2013; Muller, 1930; Nimmo et al., 1994). In contrast, theoretical work suggests that fission yeast heterochromatin displays fundamentally bistable behavior, indicating that the ‘ON’ and ‘OFF’ states are intrinsically stable (Dodd et al., 2007). Similar bistable behavior has also been experimentally observed in plants (Angel et al., 2011; Angel et al., 2015).

To reconstruct how heterochromatin domains reform by spreading with high fidelity, and thereby maintain cell identity, we have to determine both the intrinsic capacity of the spreading reaction *in vivo* and the cellular pathways that may exist to tune its behavior. The simplified heterochromatin systems in yeasts, especially *S. pombe*, are ideal for dissecting how heterochromatin spreading can form and maintain epigenetic states at loci critical to specifying cellular identity. In this work, we utilized the *S. pombe* system to examine heterochromatin formation with a “heterochromatin spreading sensor” (HSS), which examines spreading separately from nucleation. This advances prior efforts, where only nucleation-proximal silencing events were detected with one or two reporters (Bintu et al., 2016; Hathaway et al., 2012; Obersriebnig et al., 2016; Osborne et al., 2009; Xu et al., 2006). Our system enables precise, quantitative, and specific documentation of both nucleation and spreading reactions in single cells of *S. pombe*, allowing us to monitor the intrinsic behaviors of both reactions. Using the HSS, we show that different nucleators trigger distinct classes of spreading, and collaborate to form a high fidelity domain. The type of strategy we uncover has important implications for how heterochromatin spreading achieves and maintains “epigenetic” character and can safeguard cell identity against environmental perturbations.

## RESULTS

### A. A single cell heterochromatin spreading sensor (HSS) controls for nucleation and cellular noise

To assess the intrinsic behavior of heterochromatin spreading and its fidelity, we employed transcriptionally encoded fluorescent reporters to read silencing by heterochromatin at a given locus, as previously reported. Several critical improvements over prior systems enable documentation of the spreading reaction at high sensitivity (Bintu et al., 2016; Hathaway et al., 2012; Obersriebnig et al., 2016; Osborne et al., 2009; Xu et al., 2006). First, our system has high signal to noise and minimized delay from epigenetic changes to fluorescent output. We accomplish this using the weak, well-characterized *ade6* gene promoter (*ade6p*) (Allshire et al., 1994; Kagansky et al., 2009) to drive production of bright, fast-folding fluorescent proteins (XFPs) (Al-Sady et al., 2016). Second, our system provides separate sensors for nucleation, spreading, and cellular noise. We used *ade6p*-driven recoded super-folder GFP (Pedelacq et al., 2006) (“green”) and monomeric Kusabira Orange (Sakaue-Sawano et al., 2008) (“orange”) to report on nucleation and spreading, respectively (**Figure 1A**). A third XFP, *ade6p*-driven triple fusion of E2Crimson (Strack et al., 2009) (“red”, noise filter), is fully uncoupled from heterochromatin and inserted in a euchromatic locus. Here it reports on intrinsic or extrinsic noise that arises from cell-to-cell variation in the content of specific and general transcription factors and also translational efficiency (**Figure 1A**). To validate this reporter system, we characterized the non-heterochromatic state, via null mutation of *clr4* (*Δclr4*), encoding the only *S. pombe* H3K9 methyltransferase. We show that in the absence of heterochromatin, expression of the noise reporter (“red”) correlates well with that of reporters for both nucleation (“green”) and spreading (“orange”) (**Figure S1A**). Thus, cellular noise is controlled by dividing the signals from the proximal “green” and distal “orange” heterochromatic reporters by the signal of the “red”, euchromatic reporter (“green”/“red”; “orange”/“red”). Together, these elements constitute our heterochromatin spreading sensor (HSS).

### Spreading from ectopic RNAi nucleators is stochastic and produces intermediate states

To isolate heterochromatin formation from the influence of any regulatory elements that might influence the reaction, we first studied heterochromatin spreading in an ectopic context. We constructed the initial ectopic HSS based on a strain where a part of the centromeric RNAi-driven nucleation element (*dh*) is inserted proximal to the endogenous *ura4* gene (Canzio et al., 2011; Marina et al., 2013). We replaced the *ura4*+ ORF with “green” to track nucleation. To monitor spreading, the “orange” spreading sensor was inserted at one of several downstream sites from “green” (*ura4::dhHSS*^1kb^, *ura4::dh*HSS^3kb^, *ura4::dh*HSS^5kb^ *ura4::dh*HSS^7kb^, **Figure 1B**). The noise filter (“red”) was inserted between *SPBC1711.11* and *SPBC1711.12*, a *bona fide* euchromatic region (Garcia et al., 2015). All strains were initially constructed in a *Δclr4* background, and we initiated heterochromatin formation by crossing in *clr4*+. We assessed heterochromatin formation after ~ 80-100 generations by quantifying the production of “green” and “orange” as proxies for nucleation and spreading. This period is significantly longer than ~ 25 generations timeframe required for full formation of a heterochromatic domain (Obersriebnig et al.,

2016), ensuring that the population is at equilibrium.

To quantitatively assess the states produced by spreading, we performed steady-state flow cytometry on log-phase cells, which were size-gated for small, recently divided cells (~91% G2, **Figure S1B** and supplemental experimental materials) to remove size- and cell cycle-related effects. We observed that cells populate a wide range of nucleation states rather than a single state, with the distribution of repressed states varying among the HSS distance sensor strains (*ura4::dh*HSS^1-7kb^, **Figures 1C and S1C**). To specifically examine cells that have fully nucleated, we applied a computational “nucleation clamp” that isolates cells with a “green” signal that is lower than the mean plus two standard deviations of wild-type cells containing no XFPs (see Supplemental Methods). We find that spreading is stochastic in fully nucleated cells, with some cells exhibiting full repression, but others partial or no repression of the spreading reporter. The proportion of cells that are fully repressed by spreading declines linearly with distance (**Figure 1C**; compare with scheme in **Figure 1B**). Intriguingly, cells that are not fully repressed usually exhibit intermediate levels of repression, where the mean repression shifts progressively towards maximal de-repression as a function of distance in an analog manner.

We next assessed the nature of these intermediate states in the 3kb distance reporter strain, where ~30% of cells had maximal repression at the “orange” locus and the remainder had intermediate states ranging from strongly to weakly repressed. Using Fluorescent Activated Cell Sorting (FACS), we gated for successful nucleation in the “green” channel and then binned the “orange” channel for fully repressed (low), intermediate and de-repressed (high) populations (**Figure 1D**, cartoon). We queried each bin for molecular events associated with heterochromatin formation, using RT-qPCR to determine the expression levels of “orange”, and Chromatin Immunoprecipitation (ChIP) to query the presence of the marks H3K9me2 and H3K4me3. These marks are thought to be mutually exclusive, associating with repressed heterochromatin and active promoters, respectively (Noma et al., 2001). The message level of “orange” is tightly repressed in the “low” population (0.05 of max), partially repressed in the intermediate population (0.3 of max), and nearly fully “de-repressed” (0.8 of max) in the “high” population. Thus, cells with intermediate fluorescence also exhibit partial gene repression, demonstrating that these two parameters are correlated (**Figure 1D**, RT primers indicated in diagram in 1C, solid line). Histone modification levels also correlated well with the HSS signals (**Figure 1E**, ChIP primers indicated in diagram in 1C, dashed line). The “low” fluorescence population has high H3K9me2 (0.9 of *dh*, positive control) and low H3K4me3 (0.09 of actin, positive control); the intermediate population had intermediate H3K9me2 (0.49 of *dh*) and H3K4me3 (0.23 of actin), and the high population had low H3K9me2 (0.2 of *dh*) and higher H3K4me3 (0.44 of actin). Hence, successfully nucleated cells with intermediate fluorescence also exhibit intermediate amounts of the mRNA for “orange” and histone marks reflecting heterochromatin (H3K9me2) and transcriptional activity (H3K4me3). These results support the notion that intermediate states of repression observed by cytometry represent intermediate states of spreading.

These observations are not due to the particularities of the ectopic site chosen or the behavior of the XFPs, as our results are recapitulated at the *his1* locus (*his1::dh*HSS^3kb^, **Figure S1C**), which contains only one gene (*rec10*) in the “spreading zone”, rather than several transcriptional units. Additionally, switching the nucleation and spreading reporter fluorophores produced similar results (**Figure S1C**). These results suggest that RNAi-driven heterochromatin spreading is intrinsically stochastic and multimodal, producing intermediate states of repression. This behavior is not compatible with the epigenetic behavior of endogenous heterochromatin loci.

### RNAi- and Atf1/Pcr1 nucleate two types of spreading reactions at MAT

We next examined spreading behavior at the endogenous mating type locus (MAT), which has the hallmarks of a *bona fide* high-fidelity locus. This locus is very tightly repressed (Grewal and Klar, 1997; Thon et al., 2002) and is able to faithfully propagate its gene expression state even when partially disrupted (Grewal and Klar, 1996). The MAT locus has two known heterochromatin nucleators: the RNAi-dependent *cenH* element, homologous to the *dh* fragment we inserted at *ura4* and *his1*, and the RNAi-independent element termed *REIII* (Jia et al., 2004; Thon et al., 1999). At *REIII*, two stress-responsive transcription factors, Atf1 and Pcr1, which form a heterodimer (Wahls and Smith, 1994), recognize two DNA binding sites within *REIII*, and directly recruit H3K9 methylase Clr4 and Swi6/HP1 (Jia et al., 2004; Kim et al., 2004). We validated that MAT retains its well-documented tight repression following insertion of the HSS, placing the “green” reporter within the *cenH* nucleator, and the “orange” reporter proximal to the *REIII* nucleator. Both colors were fully repressed in the large majority of cells (**Figure 2A**), which is reproduced when the color orientations are reversed (**Figure S2A**). However, for both reporter configurations, the *REIII* proximal color showed a small proportion of cells that are slightly de-repressed compared to the *cenH* internal color, consistent with previous findings (Thon and Friis, 1997). We conclude that the HSS can be used to dissect spreading at the MAT locus.

We then examined spreading in cells nucleated solely by the *cenH* RNAi-element. The *REIII* nucleator was inactivated either by deleting the critical cis-acting Atf1/Pcr1 binding sites, to create a strain designated *ΔREIII*^HSS^ (**Figure 2B**), or by disrupting the *trans*-acting factor encoded by the *pcr1* gene (**Figure S2B**). Both inactivated *REIII* strains behaved similarly, but our further analysis uses only *ΔREIII*^HSS^ to avoid complications from inactivating the *pcr1*-dependent stress response (Watanabe and Yamamoto, 1996). To our surprise, given the high-fidelity character of the MAT locus, *cenH* RNAi nucleated spreading in the *ΔREIII* strain behaved almost exactly like spreading from the ectopic RNAi-driven strains. Spreading was highly stochastic, largely forming intermediate states (**Figure 2B**). The higher nucleation efficiency in *cenH* relative to the ectopic RNAi sites likely reflects the different placement of the “green” nucleation reporter, within the *cenH* nucleation element for the MAT^HSS^ strain but adjacent to the nucleation element for ectopic reporters. Thus, the two site mutations abrogating *REIII*-mediated nucleation, convert the MAT locus from a high-fidelity site to a stochastic, multimodal locus.

To examine spreading from the intact *REIII* element, we used the historical *ΔK* strain, where the entire *cenH* nucleation element is deleted and replaced with a *ura4*+ reporter (Grewal and Klar, 1996). We introduced the HSS into this context (*ΔK*^HSS^, **Figure 2C**), using the *REIII* proximal “green” reporter to detect nucleation and the distally placed “orange” reporter to detect spreading. Although *ΔK*^HSS^ has very weak nucleation compared to strains with intact RNAi elements, its distribution is sharply bimodal: Cells were either repressed (‘OFF’, lower left corner) or de-repressed (‘ON’, upper right corner; **Figure 2C**). The tightly repressed nature of the “orange” spreading reporter is even more obvious when we consider only fully nucleated cells (green^OFF^; **Figure 2C inset**), where all cells showed a Gaussian distribution around the value of maximal repression (defined similarly as for the “nucleation clamp” Supplemental Methods and **Figure 2C**). This experiment indicates that spreading from the *REIII* element is deterministic without detectable intermediate states. Interestingly, *REIII* is not capable of effective nucleation and spreading at an ectopic site (**Figure S2D**). Due to the nature of the *cenH* deletion in the *ΔK* strain, the *REII* element, which triggers silencing in an H3K9me-independent fashion (Hansen et al., 2011), is significantly closer to the “orange” color than in *ΔREIII*^HSS^. To investigate whether *REII* contributes to *ΔK*^HSS^ behavior, we replaced the *REII* element with the *Saccharomyces cerevisiae LEU2* auxotrophy gene in the *ΔK*^HSS^ strain. Importantly, spreading remains deterministic in *ΔK*^HSS^ *ΔREII::LEU2* (**Figure S2C**), suggesting that *REII* does not contribute to the *ΔK*^HSS^ spreading phenotype. We conclude that deterministic spreading is a hallmark of *REIII* at the MAT locus.

### Multi-generational single cell imaging reveals RNAi-driven spreading to be unstable

Our measurements thus far cannot reveal the dynamics of transitions between states. This requires longterm imaging of cells over a substantial number of generations (>20), which is difficult with traditional microscopy because of cell crowding effects. Here, we use the Fission Yeast Lifespan Micro-dissector (FYLM) microfluidic device (Spivey et al., 2017; Spivey et al., 2014), which traps the old pole of a rod shaped *S. pombe* cell at the bottom of a chamber well for its entire lifetime. Sibling cells generated at the new pole by medial fission eventually exit the chamber. We continuously image the old-pole cell with fluorescence microscopy for up to 60hrs (**Figure 3A**). We note that unlike *Saccharomyces cerevisiae, S. pombe* does not execute an aging program but rather dies stochastically (Coelho et al., 2013; Nakaoka and Wakamoto, 2017; Spivey et al., 2017). Thus, imaging *S. pombe* over long timescales avoids the confounding effects of aging on epigenetic behavior (Guarente, 2000; Li et al., 2017). To capture the long-range dynamics of spreading, we imaged approximately one hundred cell of each strain concurrently. Cells that maintained nucleation were analyzed further to address spreading (see **Figure S3B** for a summary of cell fates). For each cell, we imaged all three channels continuously, and performed similar normalizations as for the flow cytometry data (supplemental experimental procedures). We first imaged the HSS distance sensor strain (ectopic *ura4::dh*HSS^3kb^) (**Figure S3A**). This strain shows unstable nucleation, consistent with our flow cytometry data (**Figure 1C**). However, over time intervals where nucleation persists, we observed dynamic fluctuations in the distal “orange” color without a fixed temporal pattern (**Figure S3A** and **SVideo 1** and **2**), which is not due to the repression state of “green” (**Figure S3F**).

Next, we analyzed the MAT locus strains and selected cells that maintained nucleation for their entire measured lifespan (supplementary methods). Under this constraint, the three strains exhibit vastly different behaviors (**Figure 3B**). Wild-type MAT^HSS^ cells maintained “orange” repression for the majority of their measured lifespans (**Figure 3C, S3C** and **SVideo 3**). However, we documented transient loss of “orange” silencing for 20% of the cells. (**Figure 3B** and **3C**). In contrast, while most cells stay similarly nucleated in *ΔREIII*^HSS^ (**Figure 3D, S3D**) 83% of the cells imaged experienced at least half-maximal “orange” de-repression at some time points (**Figure 3B**). For this strain, 30% of the cells transited through the fully ON state (**Figure 3B, 3D, S3D** and **SVideo 4**). In fact, cells sampled a wide range of values from OFF to fully ON, indicating that cells do not occupy ON or OFF states exclusively, but adopt intermediate values across time (**Figure 3D**). Importantly, *ΔREIII*^HSS^ cells, just as *ura4::dh*HSS^3kb^ cells, fluctuate in their “orange” values, indicating that spreading adopts a random walk type behavior. To analyze *ΔK*^HSS^ cells, which exist predominantly in fully “green” and “orange” ON state (**Figure 2C**), we isolated OFF *ΔK*^HSS^ cells by first streaking for single OFF colonies. OFF-enriched *ΔK*^HSS^ behaved markedly differently from *ΔREIII*^HSS^: in all of the cells analyzed, “green” and “orange” reporters remained OFF throughout the whole time course (**Figure 3B, 3E, S3E** and **SVideo 5**). These data indicate that maximal spreading in these cells is fully maintained in a tight manner up to 25 generations, revealing a fundamentally different dynamic behavior in the spreading from *cenH* or *REIII*.

### Memory formation at MAT is dependent on *REIII*

To probe memory capacity (i.e., the ability of cells to retain information of an ancestral state established many generations prior) we compared cells containing an intact MAT locus to those lacking either *RNAi*- or REIII-nucleated spreading. We established two ancestral states (**Figure 4A**) with either unperturbed heterochromatin or with fully-disrupted heterochromatin using the HDAC inhibitor trichostatin A (TSA; full erasure after 10 generations of treatment, **Figure S4**). Following production of the ancestral states, we grew cells either in rich media alone or in a TSA concentration gradient for 25 generations and then measured the fraction of fully nucleated cells that effectively silence the “orange” spreading marker (**Figure 4A**). If the fraction of the population with full spreading (“orange”^OFF^) depends on the ancestral state, then cells exhibit memory. Memory is indicated by separation of the unperturbed (light orange) and perturbed (red) lines, whereas no memory is indicated by convergence of the two lines (graphs in **Figure 4B-D**). We further defined the relative “persistence” of the heterochromatin spreading as the degree to which a strain maintains “orange”^OFF^ along the TSA concentration gradient. Persistence is quantified as the TSA concentration at which the fraction of cells with “orange”^OFF^ declines to 50% of the no TSA pretreatment value (analogous to an EC^50^ value). This experimental setup allows us to directly measure the balance between history dependence, or memory, and sensitivity to perturbation.

As expected, wild-type MAT^HSS^ exhibited obvious memory at 25 generations (**Figure 4B**), which was still weakly evident even at 35 generations (**Figure S4C**). Among fully nucleated (“green”^OFF^) cells, those that derived from untreated ancestral cells showed a greater fraction of silencing (“orange”^OFF^) than those derived from treated cells throughout the entire TSA gradient, with a half-persistence point of ~2 μM (**Figure 4B**). Thus, wild-type MAT^HSS^ memory is robust in the face of perturbations of the heterochromatic state.

In sharp contrast, when spreading exclusively nucleates from RNAi (*ΔREIII*^HSS^ strain), memory of silencing (“orange” off) is significantly weaker. History dependence collapsed beyond low TSA concentrations (> 0.2 μM TSA), with the red and orange lines coinciding for much of the gradient. Even at 0 μM TSA, history dependence was erased at 35 generations (*Figure S4C*). Interestingly, the half-persistence point was ~0.2 μM, 10-fold lower than that of WT MAT (*Figure 4C*). As cenH-nucleated spreading in *REIII*^HSS^ produces little memory capacity and lacks persistence, the memory capacity at MAT therefore does not derive from RNAi-nucleated spreading.

The high persistence and memory capacity of MAT^HSS^ appears to derive specifically from *REIII* nucleated spreading (*ΔK*^HSS^ strain). This strain has a half-persistence point of ~ 3 μM TSA (**Figure 4D**), similar to the intact locus. Significantly, *REIII* nucleated spreading produces a very strong history dependence: whereas untreated ancestral cells maintained repression, ancestral cells pretreated with TSA were completely unable to repress “green” or “orange” above the *Δclr4* background even at 0 μM TSA. Remarkably, the extreme difference observed at 0 μM TSA is maintained up to about 1 μM TSA (**Figure 4D**), and does not decline at 35 generations (**Figure S4C**). Together these results indicate that *REIII-* spreading possesses an extraordinary type of memory, and suggest that the history dependence at MAT is conferred by *REIII*.

### *REIII* imposes epigenetic behavior under environmental stress conditions

Resistance to physiologically relevant environmental perturbation is necessary to maintain epigenetic states in naturally changing environments. We examined heterochromatin persistence at different temperatures in wild-type MAT, and derivatives lacking either RNAi-nucleated (*ΔREIII*^HSS^) or *REIII*-nucleated spreading (*ΔK*^HSS^). Heterochromatin is significantly lost at elevated temperatures in the wild-type MAT and RNAi-mediated (*ΔREIII*^HSS^) strains, with both strains losing 50% of repression by spreading at 36°C, and almost all repression at 38°C (**Figure 5A**). This finding is consistent with relocation of RNAi nucleation factors to the cytosol at these temperatures (Woolcock et al., 2012). In contrast, spreading in the REIII-nucleated (*ΔK*^HSS^) strain is remarkably persistent, retaining 60-70% of spreading even at 40°C. Since a large number of *ΔK*^HSS^ do not nucleate (~80%, **Figure 2C**) and are removed from the analysis, this result only reflects cells that are *REIII* nucleated.

We next studied how *REIII* contributes to retaining or reforming the heterochromatin state after a transient exposure to elevated temperature (38°C, 10 doublings) followed by return to growth at 32°C (Schematic, **Figure 5B**). As expected from our steady state experiments above, *REIII*-mediated spreading (*ΔK*^HSS^ cells) is only minimally affected by the perturbation and regains full spreading rapidly (**Figure 5D, F**), whereas wild-type MAT and RNAi-nucleated (*ΔREIII*^HSS^) strains lose a significant amount of spreading (**Figures 5C, E**) and nucleation (Figure insets). Both strains regain nucleation at *cenH* rapidly (1 day after return to 32°C; **Figure S5**), but are discrepant in their kinetics of spreading restoration: the RNAi-nucleated strain (Δ*REIII*^HSS^) requires significantly more time than wild-type MAT to recover to the 32°C extent of spreading (**Figure 5F**). Indeed, compared to wild-type MAT, *ΔREIII*^HSS^ exhibits 20 hours of lag before reaching 50% of the initial state, and plot fitting reveals a half-life (t_1/2_) difference of ~22hrs, or ~9-10 generations (Figure 5F). Therefore, *REIII*-is required for efficient recovery to the fully repressed state after heat perturbation.

Together, these data suggest that a central role of the *REIII* element is to ensure that memory of the epigenetic state at MAT predominates over environmental perturbations in the wild.

## DISCUSSION

Cell identity depends on formation of a genome partitioning pattern by heterochromatin. The ability to maintain identity depends on “remembering” the positional extent of heterochromatin spreading, rather than its nucleation, since in many systems spreading is the dominant contributor to the pattern (Schultz, 1939; Schwartz et al., 2006; Wen et al., 2009). Yet, how intergenerational fidelity that is required for memory is linked to the intrinsic properties of the spreading reaction itself has remained opaque. Surprisingly, by directly probing the fidelity of spreading in *S. pombe* with single cell assays, we found that RNAi-nucleated and REIII-nucleated spreading differ in their capacity for memory. RNAi-nucleated spreading is labile and lacks significant memory capacity despite being the most prevalent form of heterochromatin spreading and present at the MAT locus. Instead, memory formation at MAT relies on the *REIII* nucleation element, which triggers a second form of spreading that is deterministic and highly persistent. We discuss how these two qualitatively different elements at the MAT locus collaborate to sculpt a high fidelity heterochromatin locus and more broadly, how utilizing distinct forms of spreading with different epigenetic characteristics enables the organism to engineer heterochromatin elements for different biological needs.

### Different types of heterochromatin spreading exist in *S. pombe*

We have shown that *REIII* element- and RNAi-nucleated heterochromatin spreading events differ. The deterministic spreading from *REIII* is highly efficient, forming no intermediate states (**Figure 2C, 3E**), always fully switching off the spreading reporter. This type of spreading correlates with extreme memory capacity, which allows faithful intergenerational propagation of spreading that is evident in both population (**Figure 4D**) and single cell tracking experiments (**Figure 3E**), and which is consistent with previously documented bistable behaviors ascribed to the overall locus (Dodd et al., 2007; Grewal and Klar, 1996).

In contrast, RNAi-mediated nucleation leads to stochastic spreading that only occurs in some cells (**Figure 1, 2B, 3D** and **S1C**), more consistent with position effect variegation at genetically disrupted systems (Elgin and Reuter, 2013; Nimmo et al., 1994). RNAi-nucleated spreading produces intermediate states (**Figure 2B, 3D, S2B** and **S3A**) with a distinct molecular signature (**Figure 1D, E**). Our single cell tracking data indicates that cells can adopt intermediate levels of fluorescence for extended periods (**Figure 3D, S3A** and **S3D**), arguing that observed intermediates may not be a result of OFF-ON oscillations, but instead represent distinct intermediate states. Possibly, intermediate states have reduced H3K9me3, which, while not required for assembly of heterochromatin structures, is required to enact gene silencing (Jih et al., 2017). Alternatively, intermediate states might represent heterochromatin with interspersed unmethylated tails, creating “gaps” that would reduce recruitment of silencing factors via chromodomain proteins (Fischer et al., 2009) and disrupt Swi6/HP1’s oligomerization (Canzio et al., 2011; Canzio et al., 2013), resulting in lowered nucleosome stability (Yamane et al., 2011).

The proximal cause for the divergent behaviors of RNAi- and *REIII*–driven spreading is likely that they differ in stability. RNAi-spreading is very labile to both chemical (**Figure 4C**) and environmental (**Figure 5A**) perturbations, while REIII-originating structures are extraordinarily persistent under those conditions (**Figure 4D** and **5A**). We propose that high stability heterochromatin structures are more likely to undergo high-fidelity re-formation in subsequent generations, resulting in deterministic behaviors. We discuss molecular models that account for these distinct stabilities in the supplemental discussion.

### Collaboration of RNAi dependent and independent mechanisms in the formation of a high-fidelity locus

Almost all MAT^HSS^ cells faithfully propagate and remember both the spatial extent and the degree of repression at the locus as measured by our reporters, and are able to either maintain repression or quickly re-establish it after a perturbation (**Figure 2A, 3C, 4B** and **5F**). This behavior cannot be explained by *cenH* and *REIII* elements acting independently, as each alone is significantly defective in either spreading (*cenH;* **Figures 2C and 3D**) or nucleation (*REIII;* **Figure 2D** and **S2C**). Therefore, the two elements must collaborate, most likely by *cenH* stimulating *REIII* nucleation (model, **Figure 5G**). Recent findings indicate that Atf1/Pcr1 are present at *REIII* even in unsilenced, non-heterochromatic *ΔK*-type cells (Wang and Moazed, 2017). We speculate that heterochromatin originating from *cenH* stabilizes Atf1/Pcr1-dependent recruitment of silencing factors such as Clr4 (Jia et al., 2004). Given the bistable behavior of *ΔK* heterochromatin shown here (**Figure 2C** and **3E**) and in the literature (Grewal and Klar, 1996), stabilized recruitment likely becomes self-sustaining, not requiring *cenH* for maintenance.

This hypothesis is reinforced by comparing cells with the wild type MAT-locus to *REIII*-nucleated (*ΔK*^HSS^) cells during TSA recovery. Whereas *ΔK*^HSS^ cells very rarely renucleate (**Figure 4D**), *REIII* at the intact MAT locus does renucleate, as the heterochromatin reformed after erasure has much higher persistence than that nucleated from RNAi (*cenH*) (red lines in **Figure 4B** vs **C**). In contrast, we note that in continuous growth at high temperatures, heterochromatin spreading at the wildtype MAT locus resembles that of RNAi heterochromatin (**Figure 5A**). This may result from RNAi factors becoming cytosolic at high temperatures (Woolcock et al., 2012), interfering with the normal collaboration between the two elements.

While the presence of *cenH* helps raise *REIII* nucleation in steady state, *REIII* steps in under perturbation conditions to protect or quickly re-establish the heterochromatin state (**Figure 4B** and **5F**). The heat recovery experiment indicates that it is spreading controlled by *REIII*, and not effects on nucleation (**Figure S5A** vs. **B**), that take on a special role in recovery of heterochromatin lost by the collapse of *cenH* spreading (model, **Figure 5G**). Furthermore, recovery occurs 9-10 generations faster with *REIII* than without *REIII* (**Figure 5F**). *REIII* can also act to prevent the loss of heterochromatin structures in the first place, as is evident under TSA perturbation (**Figure 4B and C**). The ability to stabilize heterochromatin against perturbation by TSA, a histone deacetylase (HDAC) inhibitor, is likely due to the ability of *REIII*-bound Atf1/Pcr1 to directly recruit key HDACs (Clr3, Clr6) involved in heterochromatin formation (Kim et al., 2004; Yamada et al., 2005). We propose that in this case *REIII* increases the persistence of heterochromatin by raising the local inhibitory TSA dose.

Together, these results indicate a dynamic collaboration between two types of elements at the MAT locus to enable high fidelity, where *cenH* raises intrinsically weak *REIII* nucleation and *REIII* stabilizes or quickly re-establishes heterochromatin at the locus when *cenH* becomes compromised during perturbations (model, **Figure 5G**).

### Implications for the maintenance of heterochromatin spreading

Here we propose a model based on our data that explains why a division of labor between elements such as *cenH* and *REIII* is required for maintenance of heterochromatin spreading with high fidelity, which may extend to other systems. If heterochromatin spreading is not inherently capable of stable maintenance, auxiliary functions have to be built into heterochromatin loci to permit high fidelity. *REIII*, but not RNAi-nucleators, show indications of such auxiliary control. While RNAi nucleators behave like an autonomous element that can be transposed to any genomic context and induce spreading (**Figure 1, S1C** and **2**), *REIII* does not (**Figure S2D**, (Wang and Moazed, 2017). This inability to function outside its endogenous context points to *REIII* operating under a local chromatin structural constraint, although other models accounting for *REIII* behavior cannot be excluded (supplemental discussion). Chromatin structure formation often depends on chromosomal context as it may involve interactions with distal elements (Bonev and Cavalli, 2016; Dekker and Heard, 2015), possibly explaining the non-autonomous nature of the *REIII* memory element. Importantly, formation of folded chromatin structures has been associated with high memory in the polycomb pathway (Bantignies and Cavalli, 2011). Further, local chromatin loops have been proposed to favor memory formation (Erdel and Greene, 2016). “Memory” elements formed by adoption of specialized structures may not always be sufficient for high fidelity however, since loci capable of memory can exist in a stable ON state, as shown in *S. pombe* and plants (Angel et al., 2015; Dodd et al., 2007) (**Figure 2B**). Thus, high fidelity also requires coupling to highly efficient nucleators. The very high efficiency of RNAi-nucleators (**Figure 2A** and **2B**) render them a suitable partner. The reasons that these nucleators are not themselves capable of memory formation (at least in *S. pombe*) remain to be determined.

Data from other systems point to interactions of multiple elements enabling stable repression. In budding yeast, silencers can collaborate to stably repress a heterochromatin domain (Boscheron et al., 1996). In plants, the spreading and nucleation regions collaborate to confer epigenetic stability (Yang et al., 2017). Interestingly, in this system, unlike in *S. pombe*, it is the spreading reaction that proceeds to stabilize nucleation-induced silencing.

Organisms that feature pervasive heterochromatin, such as mammals and plants, may additionally require linking spreading to high-fidelity cellular processes, such as DNA methylation. DNA methylation is linked to DNA replication in metazoans (Arita et al., 2008) and can be directly connected to H3K9 methylation (Esteve et al., 2006; Sarraf and Stancheva, 2004). Its absence leads to destabilized and apparently stochastic H3K9 methylation in plants (Mathieu et al., 2007). However, even in mammalian systems that feature DNA methylation, intergenerational stability of heterochromatin spreading requires continuous presence of the non-enzymatic subunits of the spreading enzyme complex, including, for example, methyl histone reader proteins (Tchasovnikarova et al., 2015). Further, loci repressed by spreading are vulnerable to euchromatin invasion (Narendra et al., 2015). This implies that, even in mammalian systems, spreading is not intrinsically self-sustaining once initially triggered. Thus, in these systems linkages to DNA replication likely represent an additional level of stabilization beyond a core of collaboration between multiple elements. Overall, we propose that fidelity is not encoded in the spreading reaction, but rather can be achieved by synergistic action of separate elements, and additionally stabilized by connection to higher fidelity processes.

### Distinct forms of heterochromatin for different biological needs

We hypothesize that the ancestral form of heterochromatin served primarily in regulation of chromosome structure and genome defense, and thus is not under a tight epigenetic fidelity constraint, which likely emerged to safeguard cell identity. The majority of heterochromatin nucleation in *S. pombe* is RNAi-driven (Hansen et al., 2006) and localizes to the pericentromeres and subtelomeric regions. Pericentromeric heterochromatin, in fission yeast and metazoans, fulfills a structural and genome defense role, safeguarding proper chromosome segregation and keeping repetitive elements in check (Bernard et al., 2001; Saksouk et al., 2015). The precise role of subtelomeric heterochromatin is less clear, but given its highly repetitive nature (chromosome III) and homology across chromosomes (I and II) in *S. pombe*, it may protect against genomic instability by suppressing recombination (Cooper et al., 1997; Nimmo et al., 1998). Especially at pericentromeres, high-efficiency in establishing repression, which is intrinsic to RNAi-nucleators (**Figure 2B**), may be much more critical than high fidelity. The stochasticity of RNAi-spreading (**Figure 1C** and **2B**) is circumvented at the pericentromere by the placement of repetitive nucleators (Nakaseko et al., 1986), obviating the need for memory capacity at these sites.

Epigenetic fidelity likely arose with emergence of cell types. In simple eukaryotes, such as yeasts, cell identity specification is restricted to one site, the mating type locus. Co-expression of the silent cassettes in the repressed MAT locus, in addition to the information expressed stably from *mat1*, can result in haploid meiosis and production of low spore viability or death (Kelly et al., 1988), hence high fidelity at these sites is critical for organismal fitness. We believe this is why in addition to RNAi-nucleated spreading, the *REIII* element has also emerged at MAT.

Intergenerational repression of cell type specifier regions must be able to handle variations in the environment, so that identity is robust against perturbations. Heterochromatin spreading, unlike DNA replication, is in principle much more vulnerable to environmental changes. This is because the protein-protein and protein-DNA association and enzyme catalytic rates that define spreading vary with chemical conditions and temperature. In *Drosophila*, temperature has long been known to affect the degree of position-effect variegation, which relies on heterochromatin spreading (Chen, 1948); in *S. pombe*, temperature has been shown to affect the efficiency of RNAi-mediated nucleation (this study and Woolcock et al., 2012). We propose that specialized memory elements, such as *REIII*, evolved to safeguard cell identity loci against environmentally induced variegation.

In summary, we show that heterochromatin spreading initiated by the dominant and evolutionary conserved RNAi-elements is untethered from epigenetic capacity and is more reminiscent of other cellular polymers, such as cytoskeletal fibers. The need to tether heterochromatin to memory likely arose in evolution with the appearance of unique cell identities. Formation of epigenetic memory requires specialized and stabilized forms of spreading and auxiliary activities that exploit high fidelity cellular processes, such as DNA replication.

## ACKNOWLEDGMENTS

We thank Shiv I Grewal and Hiten D Madhani for their generous gifts of fission yeast strains. We thank Graham A Anderson and Shengya Cao for stimulating discussions, especially on hysteresis, and Brandan La for the initial Matlab scripts for cytometry data analysis. In addition, we thank Carol A Gross for substantial help with writing the manuscript and Jonathan S Weissman and Sigurd Braun for critical comments. This work was supported by grants from the National Institutes of Health (DP2GM123484) and the UCSF Program for Breakthrough Biomedical Research (partially funded by the Sandler Foundation) to B.A.-S., American Federation of Aging Research (AFAR-020) and the Welch Foundation (F-l808) to I.J.F. and the National Institute of Aging (F32 AG053051) to S.K.J. Flow Cytometry data was generated in the UCSF Parnassus Flow Cytometry Core which is supported by the Diabetes Research Center (DRC) grant, NIH P30 DK063720.

## MATERIALS AND METHODS

### Strain Construction

#### Plasmid/construct construction

Plasmids to generate constructs for genomic integration were generated by standard methods including Gibson assembly and *in vivo* recombination. *S. pombe* transformants were selected directly on dropout media for auxotrophic markers or onto rich media (YES) for 24 hours followed by selective media YES+ G418, YES+hygromycin or YES+nourseothricin).

#### Ura4 replacement method

To avoid interference of selection cassettes with heterochromatin function in our HSS, we produced “scarless” genomic integrations, lacking selection markers. To do so we marked the insertion site first with a *ura4* cassette by genomic integration and then replaced this cassette either with a XFP cassette or altered genomic sequence for site mutations. *ura4* replacements were isolated by 5-FOA counter-selection and confirmed by genomic PCR. This method was used to generate the atf/creb site deletions in PAS331, PAS332. *ura4* was targeted to the region between Mat3M and *cenH*, specifically including the two 7 base atf/creb binding sites (s1 and s2, and (Wang and Moazed, 2017)). The entire *ura4* cassette was then replaced with a construct containing the two 7 base pair deletions of s1 and s2. Point mutations and restoration of the pre-substitution locus was confirmed by PCR and sequencing.

### Flow Cytometry and FACS sorting

For standard flow cytometry experiments, cells were grown overnight in rich media (YES) and then diluted in the morning to OD=0.1 in minimal media plus supplements (EMM complete) and grown 4-6 hours before analysis by flow cytometry. Flow cytometry was performed using Fortessa X20 Dual or LSRII instruments (Becton Dickinson, San Jose, CA, U.S.A). Samples sizes ranged from ~10,000-100,000 cells depending on strain growth. Compensation was performed using cells expressing no XFPs and single color controls expressing 1 XFP each. Compensated data was used for all downstream analysis. Fluorescence was detected for each color as described (Al-Sady et al., 2016). For FACS sorting experiments, cells were grown overnight from OD=0.025 in YES and the in the morning concentrated into a smaller volume to achieve a flow rate of ~5000 events/second on the cytometer. Sorting was performed using either Aria2 or Aria3u machines (Becton Dickinson). Prior to sorting cells were strained through a 35-40 μm mesh (Corning) to reduce clogs. Sorting criteria included a gate for size (forward (FSC) and side (SSC) scatter), removal of doublets, a gate for “green”^OFF^ (“green” signal within the range of an unstained control and then gated into Low, Intermediate, High “orange” signal defined by the following: Low encompassed signal overlapping that of an unstained control and High encompassed signal overlapping that of the *Δclr4* no heterochromatin control strain PAS355. Intermediate gate was set in between Low and High with about 100 fluorescence units of a gap (representing ~2% of the full range of captured fluorescence) to ensure reliable separation. The entire range of fluorescence detected was ~2.5 orders of magnitude. At least 8x10^6^ cells were collected for each population for Chromatin Immunoprecipitation and 2x10^6^ cells for RT-qPCR. Immediately after sorting, the final populations were subjected to the appropriate treatment for either Chromatin Immunoprecipitation or RT-qPCR.

### Sytox Green Staining and Cell Cycle Analysis

Cell cycle analyses were performed essentially as described (Knutsen et al., 2011). Briefly, cells were fixed with 70% ethanol, washed with 20 mM EDTA pH 8.0, and treated with RNaseA for 3 hours at 37°C. Immediately before analysis by flow cytometry, 2 μM Sytox Green (Invitrogen) in 20mM EDTA pH 8.0 was used to resuspend pelleted cells. Cells were excited with a 488 nm laser and FSC-A, SSC-A, Sytox Green-A (Area) and Sytox Green-W (Pulse Width) data were collected. Sytox Green signal was detected with a 505 nm longpass filter and a 530/30 bandpass filter. Analysis was performed in the FlowJo Software (Tree Star Inc, Ashland, Oregon, U.S.A.). Cells were gated in FSC/SSC to isolate single, small cells. A plot of Sytox Green-W vs Sytox Green-A was generated and the fraction of cells in each cell cycle phase (G2, S, and G1+M) within the FSC/SSC gate were calculated.

### Trichostatin A (TSA) gradient experiment

Cells were taken from fresh plates, and then grown overnight with shaking (Elmi) in 96-well plates containing 150 μL YES (Day -1). The next day (Day 0), cells were diluted into YES and measured by cytometry. At the end of Day 0, cells were passaged into YES+ DMSO (0μM TSA) or YES+ 50 μM TSA overnight. The next day (Day 1), cells were diluted and grown briefly into the same pretreatment conditions and the 50 μM TSA pre-treated cells were checked for complete de-repression by flow cytometry. Complete de-repression was defined as a qualitative overlap of WT and *Δclr4* profiles, with no evidence of repression. Both 0μM and 50μM TSA pretreated cells were then diluted into a gradient of TSA of eleven two-fold dilutions from 50 μM along with a twelfth 0 μM (DMSO) point. Cells were measured after ~6hrs and then passaged into the same TSA gradient conditions to continue growth. The next day (Day 2) cells were diluted from overnight growth into the same gradient as above, measured ~ 6hrs later by flow cytometry and passaged into the same gradient again overnight. The same protocol was followed for Days 3 and 4. The full experiment was performed twice at different times (biological replicate). Given the lengthy continuous growth, contamination was occasionally observed in <1% of wells. The replicate shown was chosen based on lacking contamination.

### Heat recovery experiment

Cells were taken from fresh plates, and then grown overnight with shaking (Elmi) at either 32°C or 38°C (Day-1) in 96-well plates containing 200 μL YES medium per well. In the morning, cells were diluted into 200μL YES and grown ~6hrs at the same temperature before measurement by flow cytometry (Day 0). At the end of Day 0 all cells were all diluted again into YES and grown at 32°C. The next day (Day 1) cells were diluted from overnight growth into YES at 32°C, measured ~ 6hrs later by flow cytometry and passaged into the same temperature overnight. The same protocol was followed for Days 2, 3, and 4.

### Chromatin Immunoprecipitation (ChIP) and quantification

We found that sonication of a small number of cells such as can be collected by FACS leads to a marked increase in background signal from negative control regions that was absent when ChIP was performed with larger log phase cultures (>50x10^6^ cells). To address this, ChIP was performed on each of the sorted populations with the addition of 42 x10^6^ formaldehyde fixed cells of *S. cerevisiae* W303 strain as a carrier. Additionally, ChIP was performed on a sample of W303 alone, which only produced signal equivalent to background. Sorted populations and W303 cells were fixed and pre-processed for ChIP separately, then mixed together immediately prior to lysis. Cells were crosslinked and lysates prepared for ChIP as described (Canzio et al., 2011) with the following exceptions: After lysis, the chromatin fraction was resuspended in 350μL lysis buffer and sonication performed using a Diagenode Bioruptor Pico machine at 4°C, with 16 rounds of 30 seconds ON, 30 seconds rest. ChIP was essentially as described, with the total lysate split into 4 equal technical replicate samples (after ~8% set aside as input fraction) and ChIP performed in 800 μL per sample. For two replicate samples 1 μL of anti-H3K9me2 (Abcam ab1220) antibody or 1 μL of anti-H3K4me3 (Active Motif 39159) antibody was added and the sample agitated on a Nutator overnight at 4°C. Immune complexes were collected for 3 hrs with 15 μL washed protein A Dynabead slurry (Invitrogen). Washing and downstream processing steps were essentially as described, except the “wash buffer” wash was performed once. Samples were purified using a Machery-Nagel PCR purification kit and NTB buffer for SDS containing samples. DNAs were quantified by RT-qPCR (see below).

H3K9me2 and H3K4me3 enrichments were calculated as follows: IP/input values for amplicons of interest were calculated for technical triplicates and normalized to the IP/Input values for positive controls for each antibody, *dh* for H3K9me2 and the actin promoter for H3K4me3.

### RNA Extraction and mRNA quantification

After sorting, samples were spun at 5000xg, supernatant decanted and pellets flash frozen in liquid nitrogen and stored at -80°C. For the *Δclr4* strain PAS335, cells were grown into log phase and then cell pellets were isolated in the same fashion. Total RNA was extracted in technical duplicates from the same cell pellets using the “MasterPure-Yeast RNA Purification Kit” (Epicentre), including a 30 minute DNAse treatment step post-RNA isolation. Reverse Transcription was performed with SuperScript III RT (Invitrogen), using the supplied protocol and 1.5-2μg of RNA and an oligo dT primer. Following cDNA synthesis the reaction was treated with RNAse H (New England Biolabs). cDNA samples were quantified by RT-qPCR in technical triplicates. For each sorted sample mKO2 cDNA values were normalized to actin and then divided by the max value calculated similarly from PAS355 (*Δclr4*).

### RT-qPCR

Real time quantitative PCR was performed using a BioRad CFX-384 machine. 15μL reactions were prepared, each containing 7.5μL of Applied Biosystems SYBR Select Master Mix, 4.5μL 3.3M betaine, 1.2μL of 2.5μM oligo mix, 0.8μL water, and 1μL template. The thermocycler protocol was: 2min at 50°C then 2min at 95°C followed by 40 cycles of 15sec at 95°C and then 1min at 60°C followed by a plate read. Lastly a melt curve was generated. Standards were generated with 5 fold dilutions of genomic DNA containing templates for all PCR products.

### Single-cell Microscopy

Single cells of strains PAS 387, 389, 391 and 244 (see strain table; E2Crimson under *act1* promoter) were captured in microfluidic devices as described (Spivey et al., 2017). Multi-channel fission yeast lifespan microdissectors (multFYLM) contained six independent devices (channels), each of which is capable of capturing up to 392 cells. In brief, the devices were cast in polydimethylsiloxane (PDMS, Sylgard 184, Dow Corning) using conventional soft lithography methods. Master structures were fabricated from P-doped silicon wafers (ID#452, University Wafers) and SU-8 photoresists 3005 and 2010 (Microchem, Westborough, MA). MultFYLMs were cleaned and adhered to glass coverslips (48 x 65 mm #1, Gold Seal), and then connected to syringes (60 mL, Becton-Dickson) containing YES 225 liquid media (Sunrise Science) via PFA tubing and microfluidic fittings (IDEX Health and Science). The multFYLM was maintained at 30°C in a custom staged-mounted environmental chamber on an inverted microscope (Eclipse Ti, Nikon) equipped with NIS Elements software (Nikon), a 60X air objective (CFI Plan Apo λ, 0.95 NA, Nikon) fitted with an objective heater (Bioptechs), a motorized stage (Proscan III, Prior), and an active feedback-based focusing system (Perfect Focus System, Nikon). An LED lamp (Sola II, Lumencorp) and a scientific-grade CMOS camera (Zyla 5.5, Andor) were used for fluorescent imaging. Multi-color fluorescent imaging of sfGFP, mKO2 and E2Crimson fluorophores was carried out by alternating between three filter sets mounted in a computer-controlled filter ring (Chroma 49002, 49010 and 49015, respectively). To help with the semi-automated cell identification, each channel was imaged every ten minutes via brightfield imaging (100 ms exposure, both in focus and 4 μm below the focal plane). Fluorescent images of each of the three fluorophores were taken every thirty minutes (150 ms exposure). This illumination scheme was well below the phototoxicity limit, as described previously (Al-Sady et al., 2016). Raw images were saved as uncompressed 16 bit ND2 files and further analyzed using a custom-written image analysis pipeline (see below).

Cells were grown overnight (30°C with 225 rpm shaking) to saturation in YES media, then diluted in YES to an optical density at 600 nm (OD_600_) of 0.1 and allowed to grow for approximately 5 hours to reach an OD_600_ of 0.5. Cells (60 μL at OD 0.5 in YES+2% Bovine Serum Albumin, BSA) were loaded at the entry port of the multFYLM. After cells entered individual channels, media lines were reattached and YES media was pumped through on a pulse cycle (14 min: 5 μLmin^-1^, 1 min: 55 μLmin^-1^) for the entire experiment. This flow regime was optimized to flush out occasional cell clumps that grew at the device inlets and other fluidic interfaces. Four genotypes were imaged simultaneously for 60 hours in each channel of a multFYLM device to ensure identical imaging and growth conditions. In all cases, we only analyze the innermost cell, which was the oldest cell pole (see below). Cells that were ejected or died within the first 12 hours after loading were not included in the downstream analysis.

### Single-cell image analysis

Single-cell imaging data was processed using an updated version of the custom-written FYLM Critic analysis package (Spivey et al., 2017). The source-code is available via GitHub (https://github.com/finkelstenlab/fylm). FYLM Critic performs the following automated processing on the raw images: (1) rotation; (2) jitter removal via a cross-correlation algorithm; and (3) generation of kymograph and individual cell images. The latter were used to create videos of individual cells in Fiji (Schindelin et al., 2012). The final outputs of FYLM critic are the position and contour of each dividing cell, as well as the time-dependent fluorescence intensities for each cell. These fluorescence intensities are obtained by averaging the intensity across all pixels that fall within the cell volume, as defined by the bright field images. This normalization also ensures that the fluorescence intensity is corrected for the size of the rapidly dividing cells. Time-dependent fluorescent intensities were analyzed via custom-written MATLAB scripts (version 2017a Mathworks, available upon request). Background fluorescence from the PDMS device was subtracted using catch tubes that did not receive a cell. The maximum heterochromatin reporter (GFP, mKO2) fluorescence intensity was calculated using *Δclr4* cells in the same reporter construct background. To control for expression variation across the cell cycle, the fluorescence from heterochromatin reporters was also reported as a ratio of the control fluorophore, E2Crimson. Similarly, cells fluorescing in the clamp channel were removed from analysis for MAT locus derived strains (see supplemental methods).

Single cell images generated by the FYLM Critic analysis were compiled into stacked movies using Fiji. Images in bright field and for each color channel were processed separately in batch and then later combined into a vertical stack. For each channel, 0.2% of pixels were allowed to become saturated and pixel values were normalized to the maximum range for the whole sequence in that channel. For bright field, every third image was included to match the imaging frequency of the fluorescent channels. Movies were edited for length to only include contiguous imaging sequences without loss of focus and for size to remove non-cellular debris and cells from the opposite side of the channel that entered the field of view.

After combining all color channels and bright field, the brightness and contrast were increased for cell 407 to match the red channel brightness of the other strains. Image sequences were saved as uncompressed .avi files with a rate of 15 frames per second.

**Table 1:**
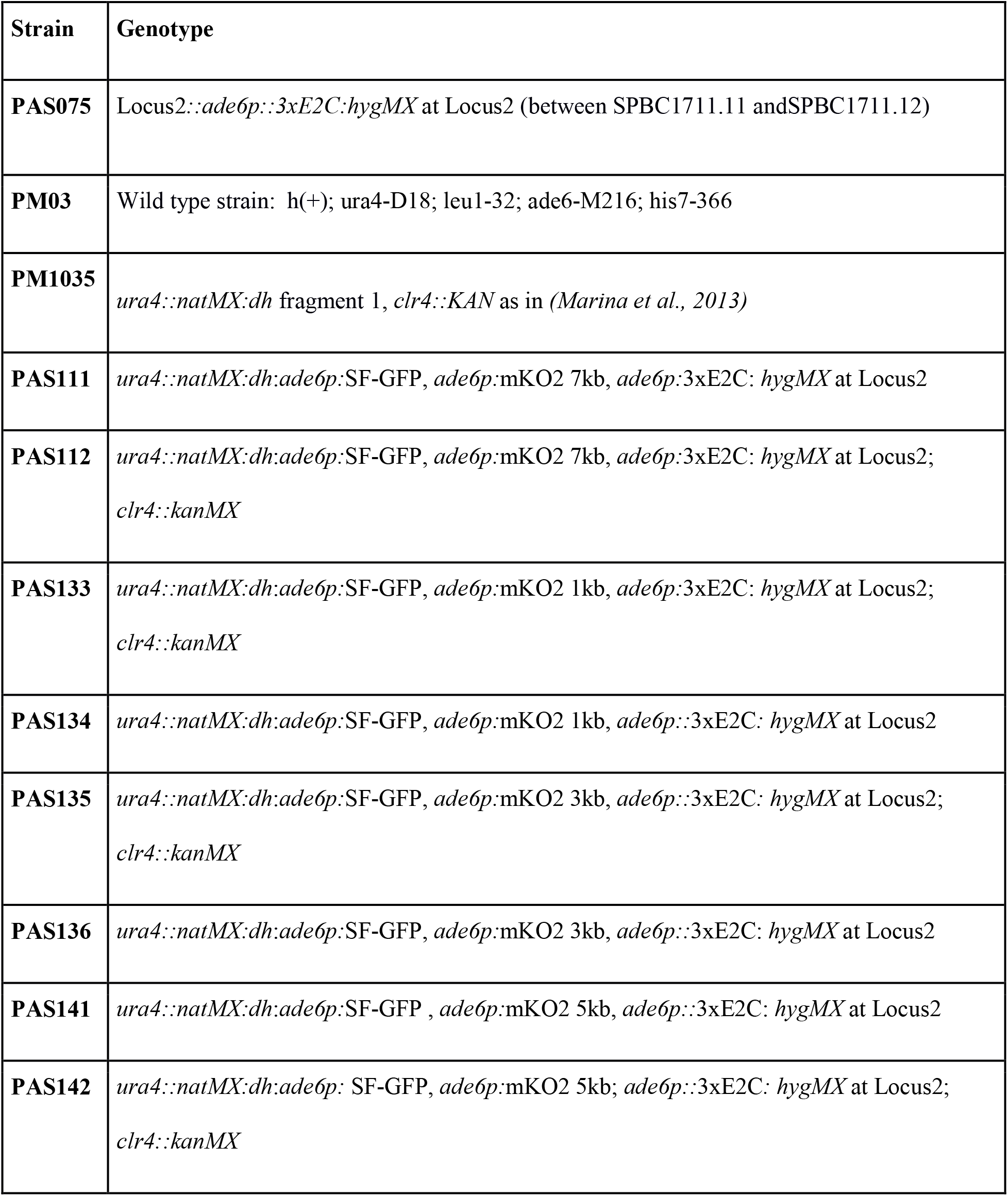
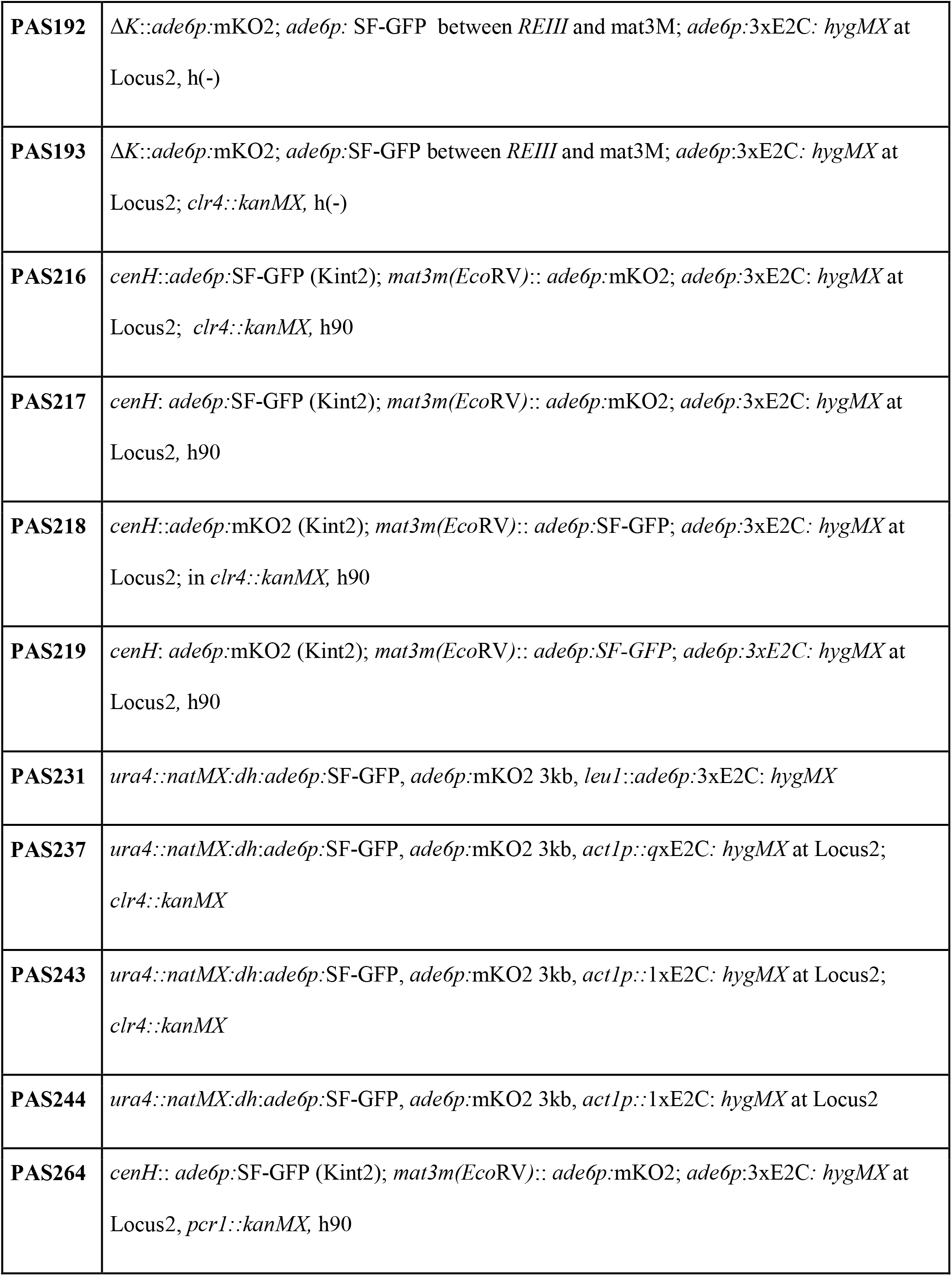
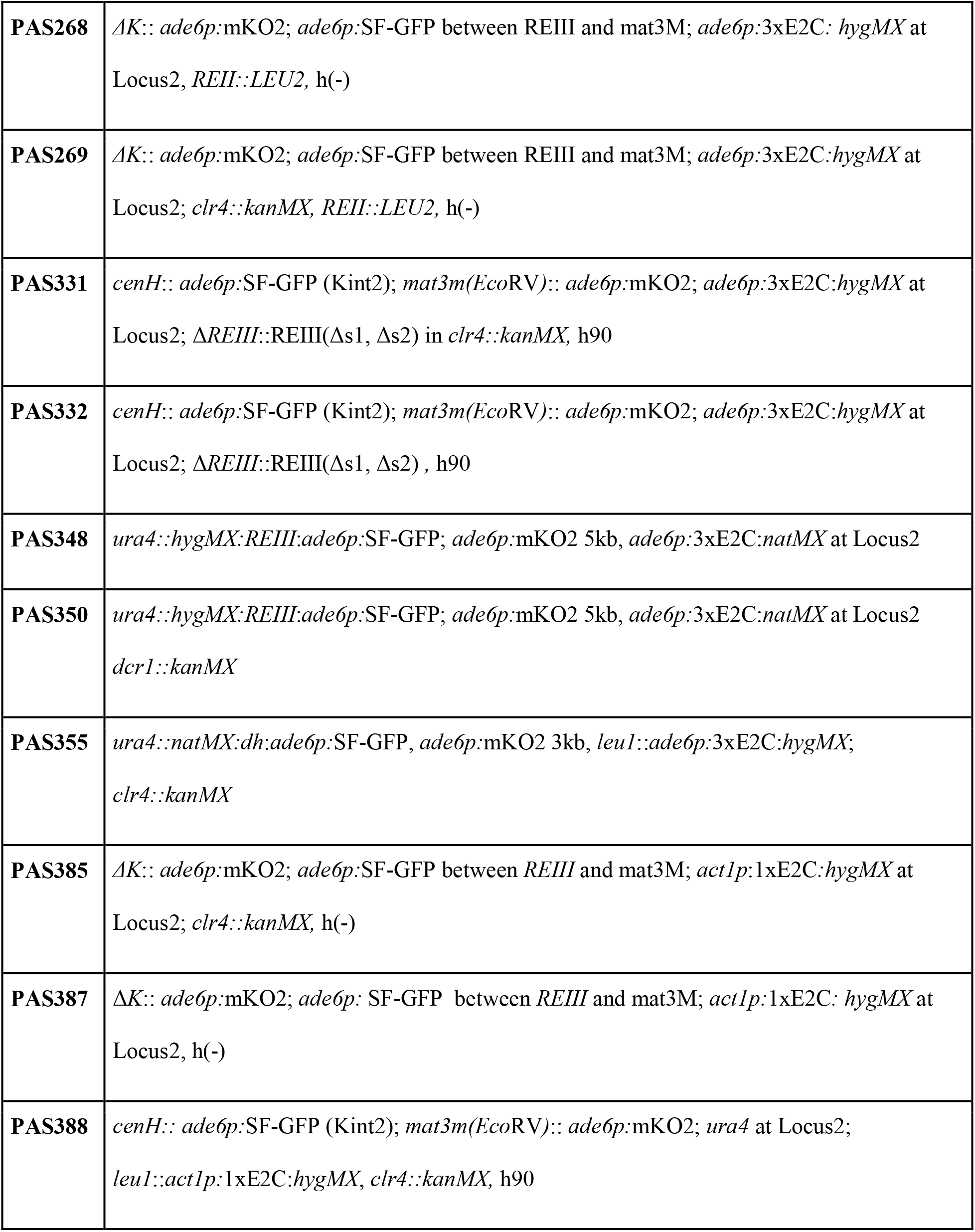
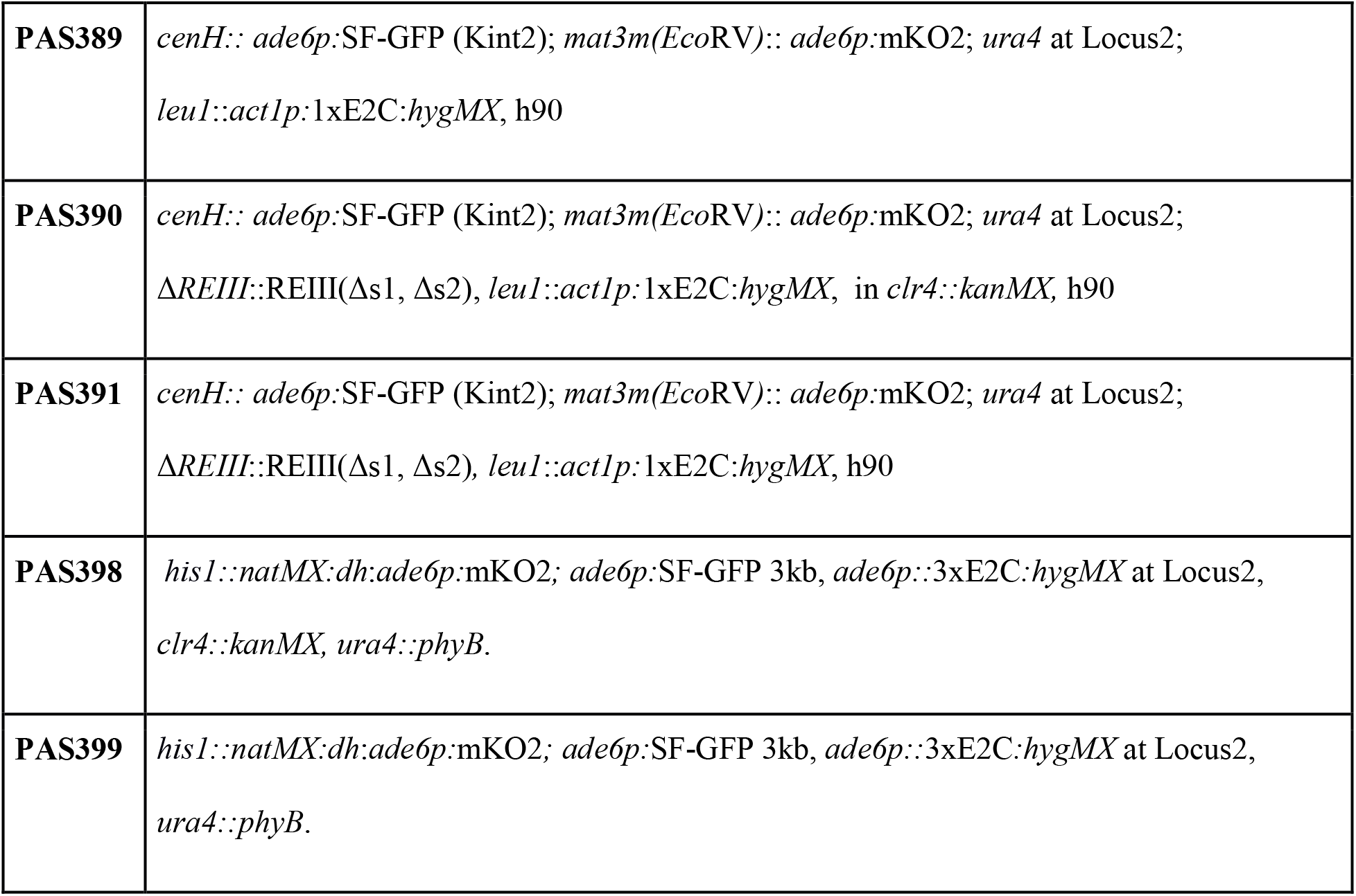
Yeast strains used in this study

**Figure 1:**
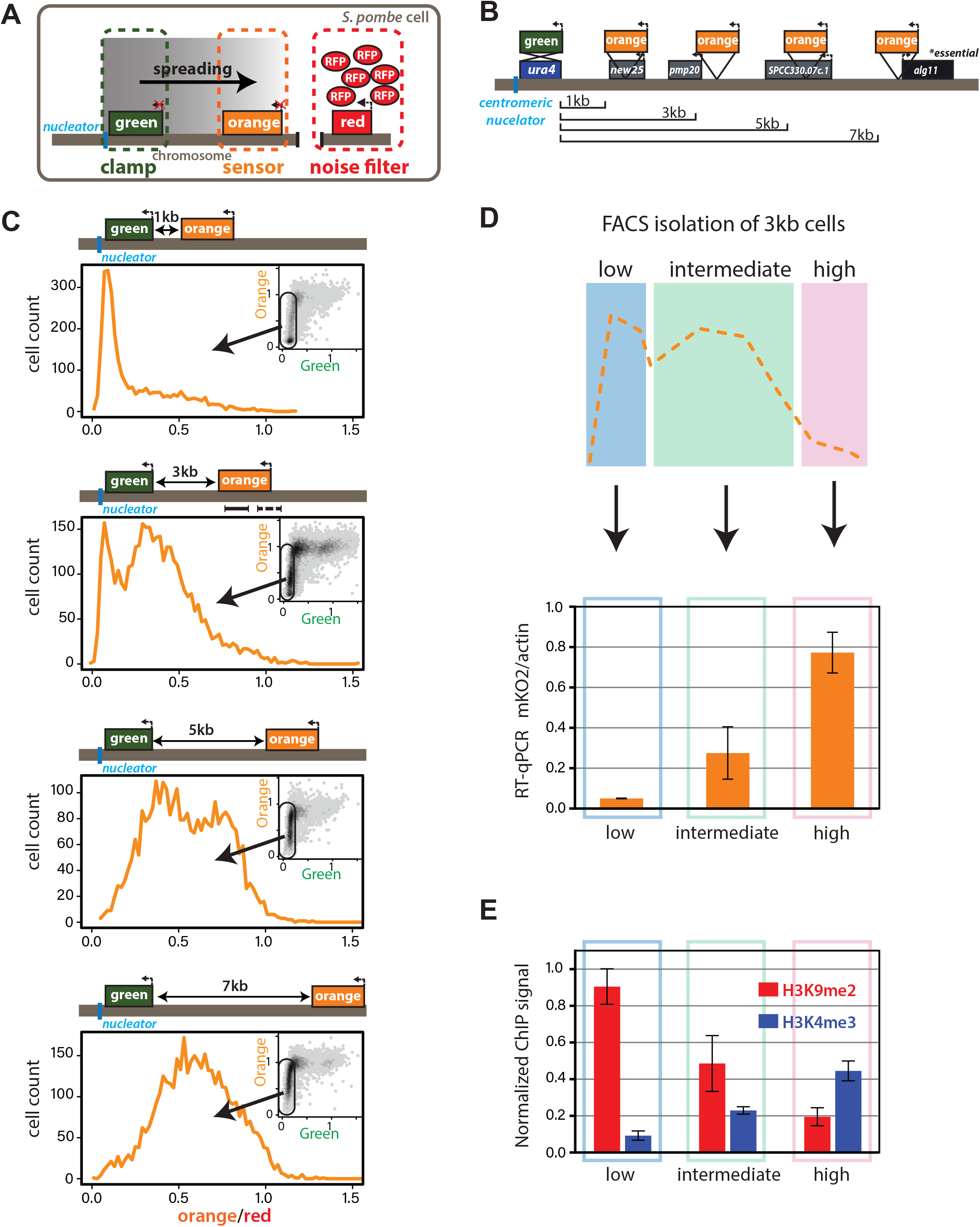
Heterochromatin spreading from RNAi-nucleated elements is stochastic and produces intermediate states. **A.** Overview of heterochromatin spreading sensor. Three transcriptionally encoded fluorescent proteins are inserted in the genome: The “clamp” site enables isolation of successful nucleation events, the “sensor” reports on spreading events and the “noise filter” normalizes for cell-to-cell noise. **B.** Overview of the *ura4::dh*HSS^1-7kb^ strains. Genes downstream of the “green” nucleation color are annotated. **C.** Spreading from *ura4::dh* visualized by the HSS with “orange” inserted at different distances shown in B. The “red”-normalized “orange” fluorescence distribution of “green”^OFF^ cells plotted on a histogram. Inset: 2D density hexbin plot showing red-normalized “green” and “orange” fluorescence within the size gate, with no “green” or “orange” filtering. The “green”^OFF^ population is schematically circled. The x-axis is normalized to =1 for the *Δclr4* derivate of each strain. **D.** TOP: cartoon overview of the FACS experiment for D. and E. “green”^OFF^ cells collected from the *ura4::dh*HSS^3kb^ were separated in three populations (“Low”, “Intermediate” and “High”) as shown schematically based on the “orange” fluorescence. BOTTOM: “orange” RT-qPCR signal for the indicated populations. The y-axis is scaled to =1 based on the “orange” signal in *Δclr4*. Error bars indicate standard deviation of two replicate RNA isolations. **E.** ChIP for H3K9me2 and H3K4me3 in the same populations as D. Each ChIP is normalized over input and scaled to =1 for a positive control locus (*dh* repeat for H3K9me2 and *act1* promoter for H3K4me3). Error bars indicate standard deviation of two technical ChIP replicates. Primer pairs for RT-qPCR and ChIP are indicated by solid and dashed line, respectively, in the C. *ura4::dh*HSS^3kb^ diagram.

**Figure 2:**
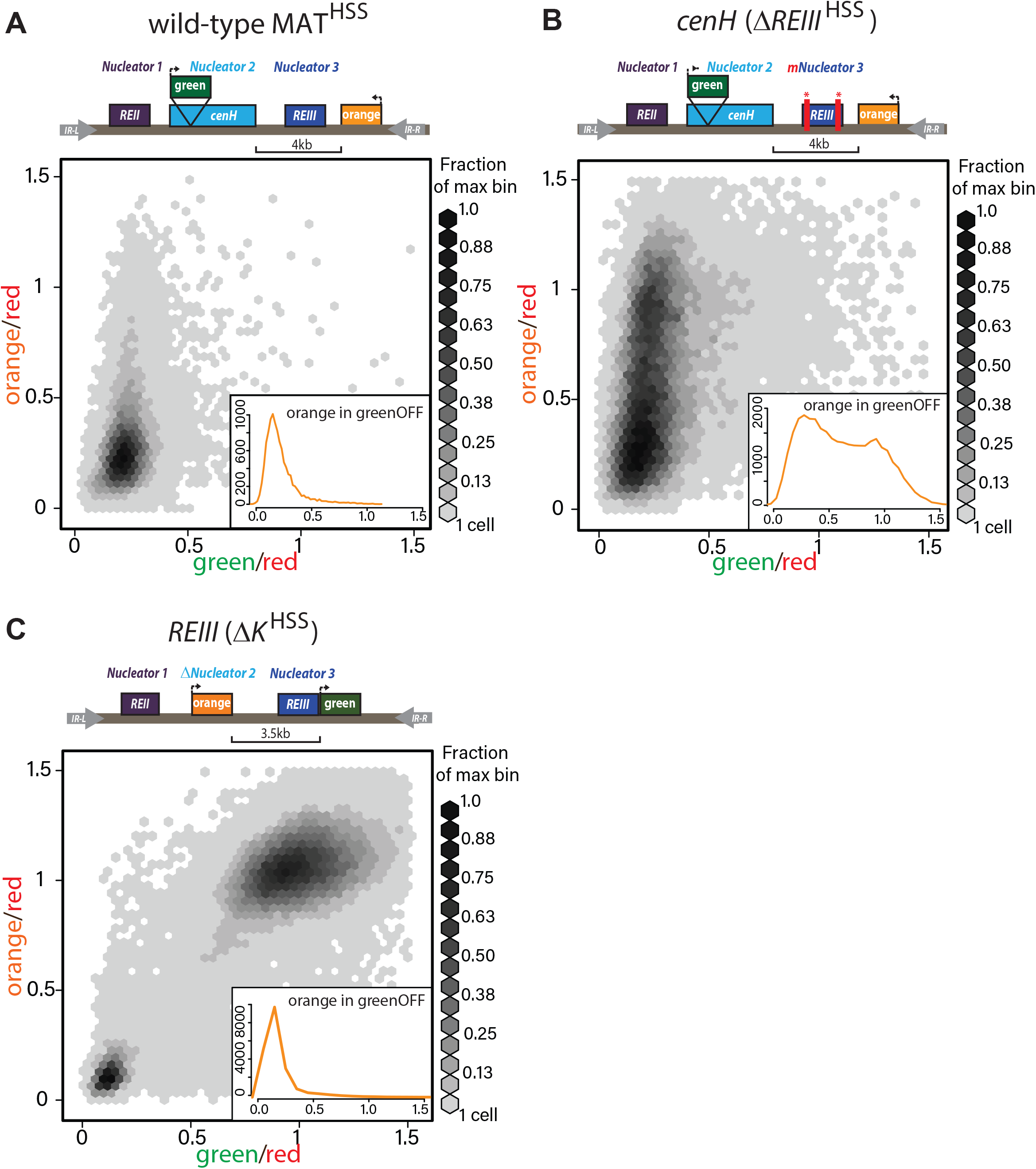
RNAi- and *REIII-* nucleators trigger qualitatively different spreading reactions in the MAT locus. **A.** TOP: diagram of the reporters within MAT^HSS^. BOTTOM: 2D-density hexbin plot showing the “red”-normalized “green” and “orange” fluorescence for wild-type MAT^HSS^ cells. Scale bar shows every other bin cutoff as a fraction of the bin with the most cells. The smallest bin starts at one cell and the maximal bin contains 498-531 cells. Inset: histogram of the “red”-normalized “orange” fluorescence distribution of “green”^OFF^ cells. B. TOP: diagram of the reporters within *ΔREIII*^HSS^, which contains two 7bp Atf1/Pcr1 binding site deletions within the *REIII* nucleation element. BOTTOM: 2D-density hexbin plot and inset as above, where the maximal bin contains 200-213 cells. **C.** TOP: diagram of the reporters within *ΔK*^HSS^. The *cenH* nucelator and additional 5’ sequence is deleted and replaced by “orange”. “green” is located directly proximal to *REIII* and serves as the nucleation clamp. 2D-density hexbin plot and inset as above, where the maximal bin contains 773-824 cells. The *Δclr4* derivative of each strain was used to normalize the X- and Y-axes to =1.

**Figure 3:**
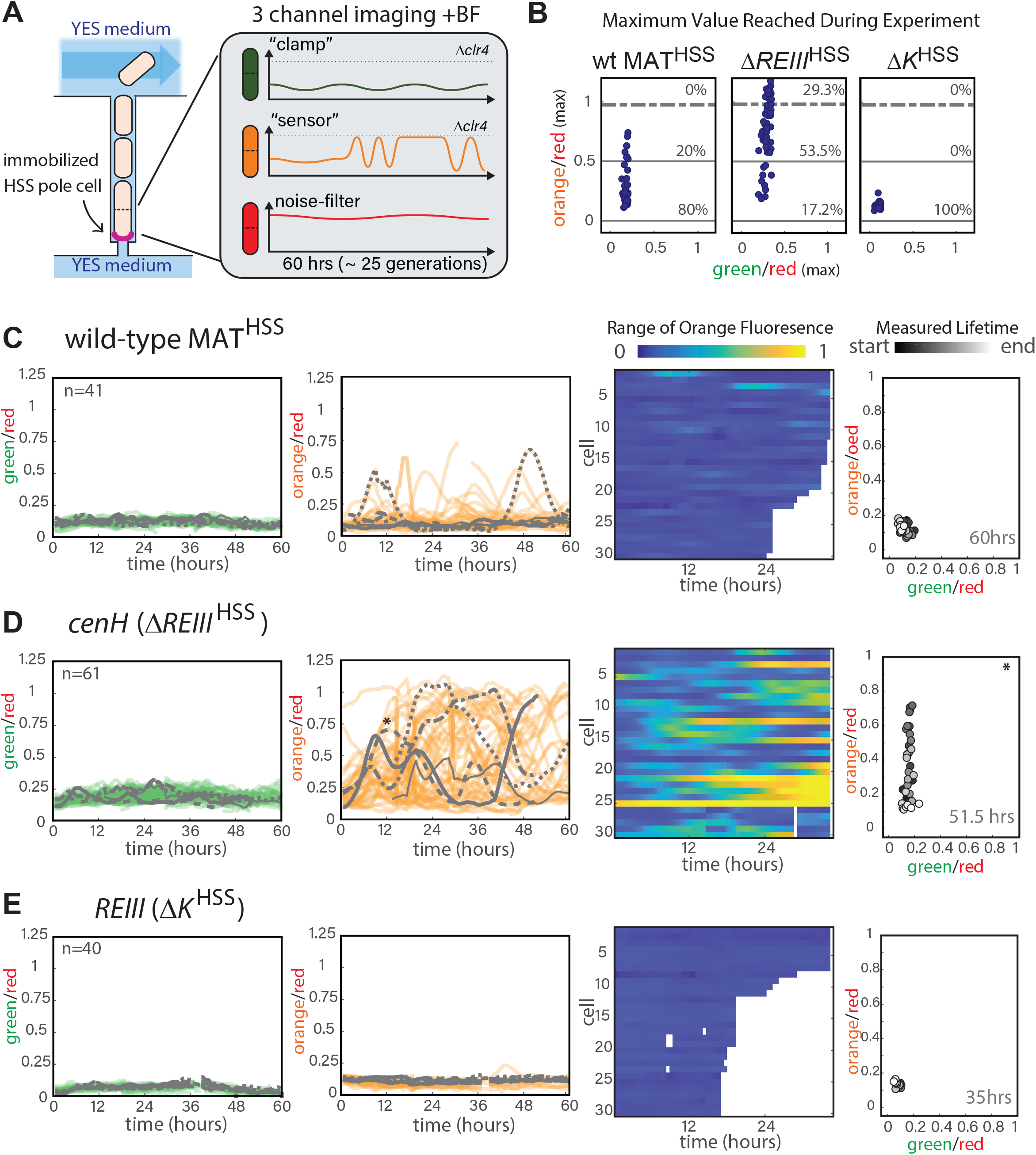
Single cell analysis of nucleation and spreading using a Fission Yeast Lifespan Microdissector (FYLM). **A.** Overview of the FYLM-based heterochromatin spreading assay. The old-pole cell is trapped at the bottom of one of hundreds of wells in the FYLM microfluidic device and is continuously imaged in brightfield (to enable cell annotation), green, orange and red channels. Hypothetical example traces are shown. **B.** Maximum values attained by each nucleated cell for normalized “orange” plotted against normalized “green”. Solid horizontal lines correspond to y=0 and y=0.5. Dashed line corresponds to an ON cutoff determined by mean less 3 standard deviations for each strain’s matched *Δclr4* strain. Percentage of cells between each line was calculated. **C.** FYLM analysis of wild type MAT^HSS^ cells. CELL TRACES: 60hrs of normalized “green” (left) and “orange” (right) fluorescence in cells that maintained nucleation with the same 5 cells overlaid in different gray line styles in both plots. Gaps indicate loss of focus. HEATMAP: Up to 36 hours of normalized “orange” fluorescence for 30 cells that maintained nucleation is represented from blue (0) to yellow (1). X-Y FLUORESCENCE PLOT: for one representative sample cell, plot of normalized “green” and “orange” fluorescence across its measured lifetime (grayscale). **D.** FYLM analysis of *ΔREIII*^HSS^ cells as in C. The example cell in the kymograph is marked with an asterisk(*) on the orange traces **E.** FYLM analysis of *ΔK*^HSS^ as in C., D. All cells were normalized to *Δclr4* (max, 1).

**Figure 4:**
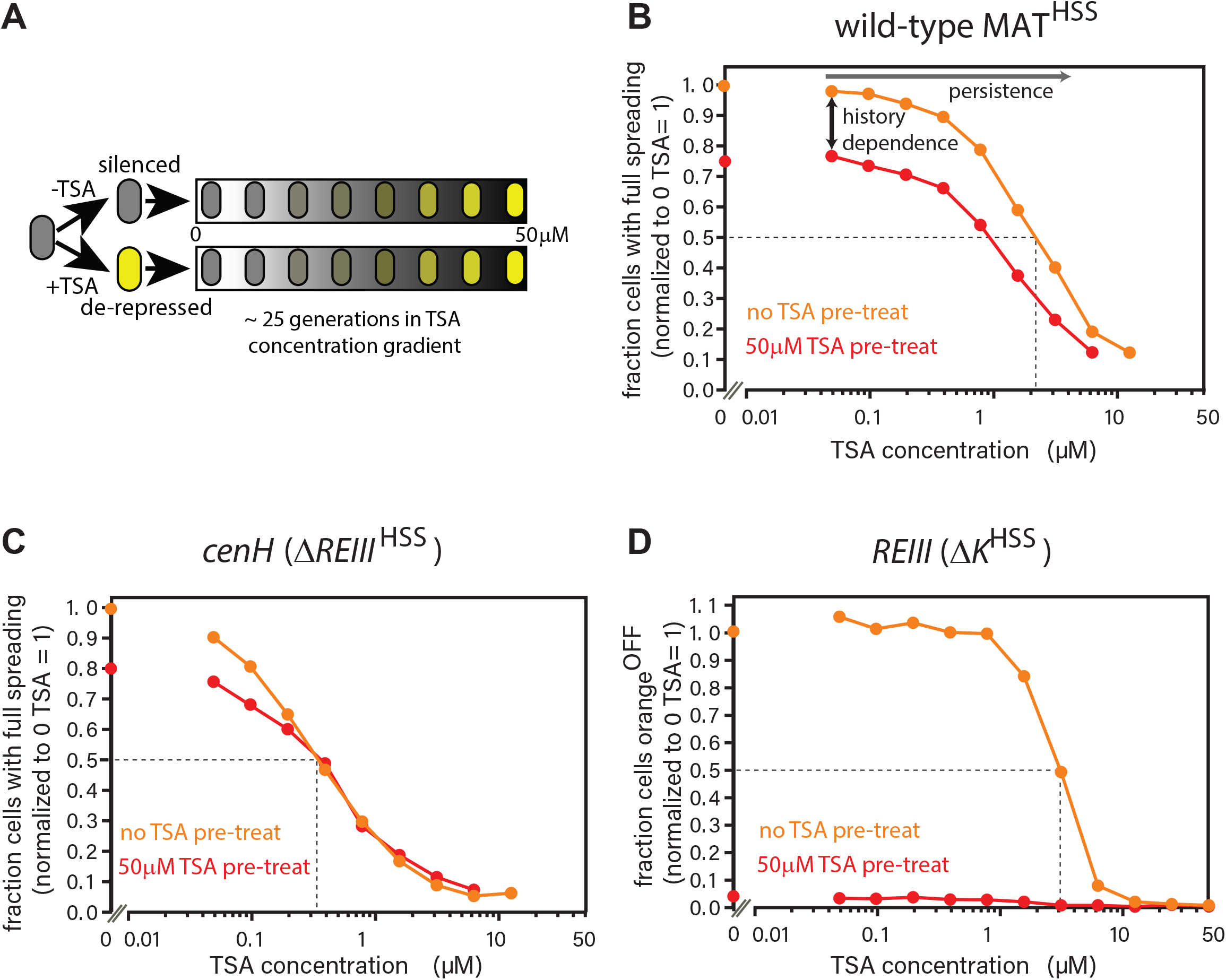
RNAi-nucleated spreading exhibits low history dependence and weak persistence compared to *REIII*. **A.** Experimental schematic for history dependence and persistence. Cells in log phase were treated with TSA (50 μM) for 10 generations to erase all heterochromatin (de-repressed, yellow) or kept untreated (repressed, gray). Both populations are then grown in a gradient of TSA concentration from 0 to 50 μM for 25 generations. **B.** The wild-type MAT locus exhibits history dependence in silencing “orange” throughout the TSA gradient. The fraction of “green”^OFF^ cells that fully silence “orange” normalized to the no TSA pre-treatment, 0 μM TSA point are plotted for each TSA concentration. Red line: cell ancestrally TSA pre-treated; Orange line: cells without pre-treatment. **C.** Spreading from *cenH* exhibits weak history dependence and low persistence. Cell populations as above. **D.** Spreading from *REIII* exhibits extreme history dependence and high persistence. The fraction of “orange”^OFF^ for all cells is plotted, because in the TSA pre-treatment almost no “green”^OFF^ cells can be detected. As spreading is deterministic for *REIII*, all “orange”^OFF^ cells are also “green”^OFF^. Dotted lines indicate the half-persistence points:TSA concentration at which 50% of non-pretreated cells fail to spread heterochromatin to “orange”. History dependence is the difference between orange and red lines. One of two full biological repeats of the experiment is shown.

**Figure 5:**
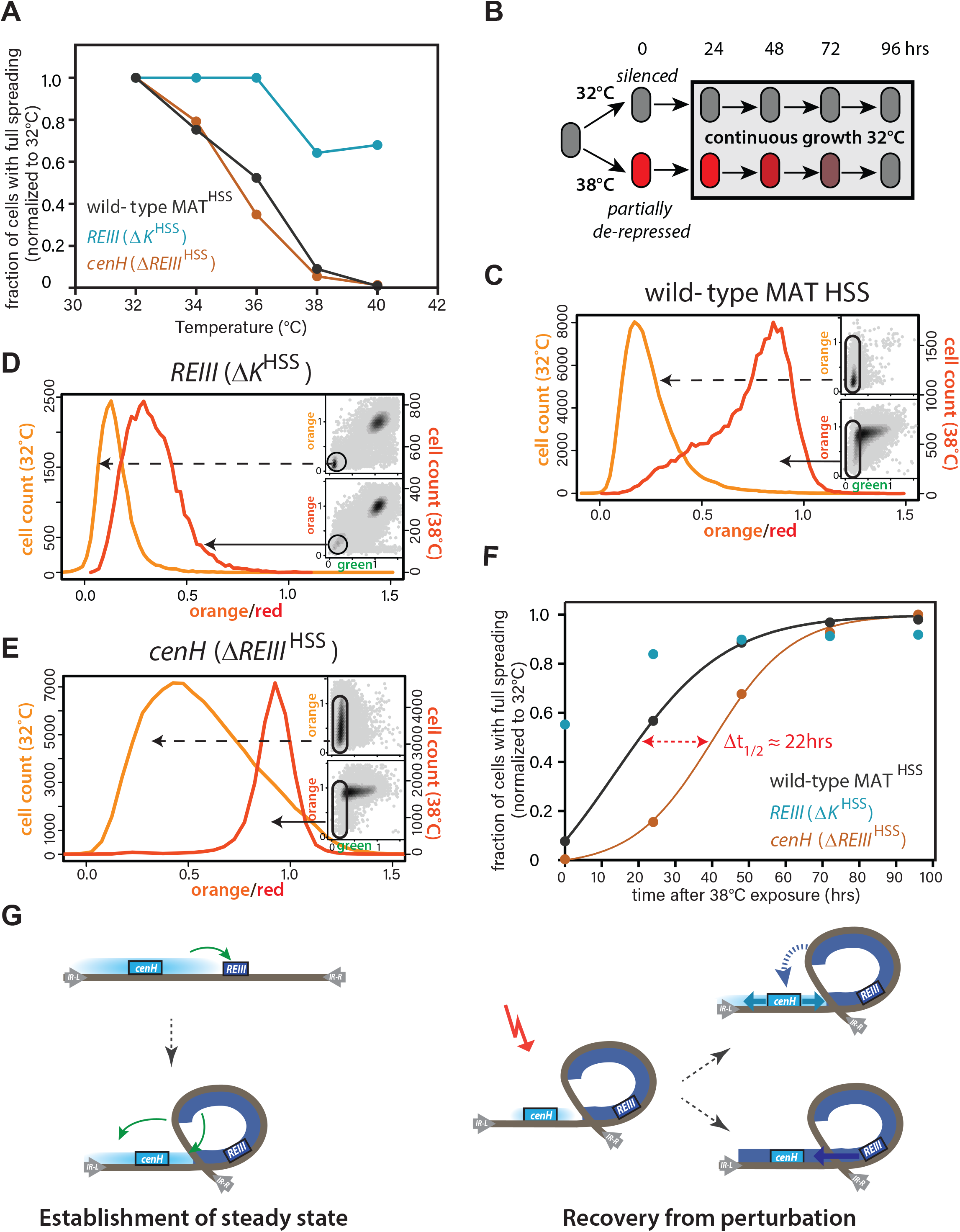
*REIII* element resists environmental perturbation and facilitates epigenetic memory by enabling rapid reacquisition of pre-perturbation state. **A.** The persistence of the heterochromatin state from 32-40°C in wild-type MAT locus^HSS^, *ΔK*^HSS^, and *ΔREIII*^HSS^. The fraction of cells that fully repress both “orange” and “green” (full spreading) at each temperature is plotted normalized to the given strains value at 32°C. **B.** Experimental schematic for heat stress and recovery. Cells were grown at either 32°C or 38°C for 10 generations and strains subsequently grown continuously for 96 hours at 32°C. **C-E.** Histograms of “red”-normalized “orange” fluorescence distribution in “green”^OFF^ cells are shown for cells grown at both 32°C (light orange) and 38°C (dark orange). Insets: 2D density hexbin plots, “green”^OFF^ cells are schematically circled. C.-E. represent t=0 in panel F. **F.** The fraction of cells with full spreading after 38°C exposure and recovery normalized to the fraction of cells with full spreading at 32°C for each strain is plotted over time. Full spreading is defined as in A. For wild-type MAT locus^HSS^ and *ΔREIII*^HSS^ strains, we fit a simple sigmoidal dose response curve and determined a t_1/2_ value. The difference in t_1/2_ values or Δt_1/2_ is ~22hrs or ~9 generations. **G.** Model for collaboration of *cenH* and *REIII* in establishing and maintaining the high fidelity MAT locus. (LEFT, TOP) During initial establishment, *cenH* heterochromatin raises the nucleation frequency at *REIII* (green arrow). (LEFT, BOTTOM). This heterochromatin may then adopt a specialized chromatin structure (indicated by looping, other models remain possible). *REIII* heterochromatin stabilizes heterochromatin at MAT (green arrows). (RIGHT) A perturbation disrupts labile *cenH-* nucleated spreading. *REIII* assists in reestablishment of the initial state either by accelerating spreading from *cenH* (blue dashed arrow, TOP), or expanding heterochromatin spreading from *REIII* (BOTTOM).

---

**Supplemental Figure 1:**
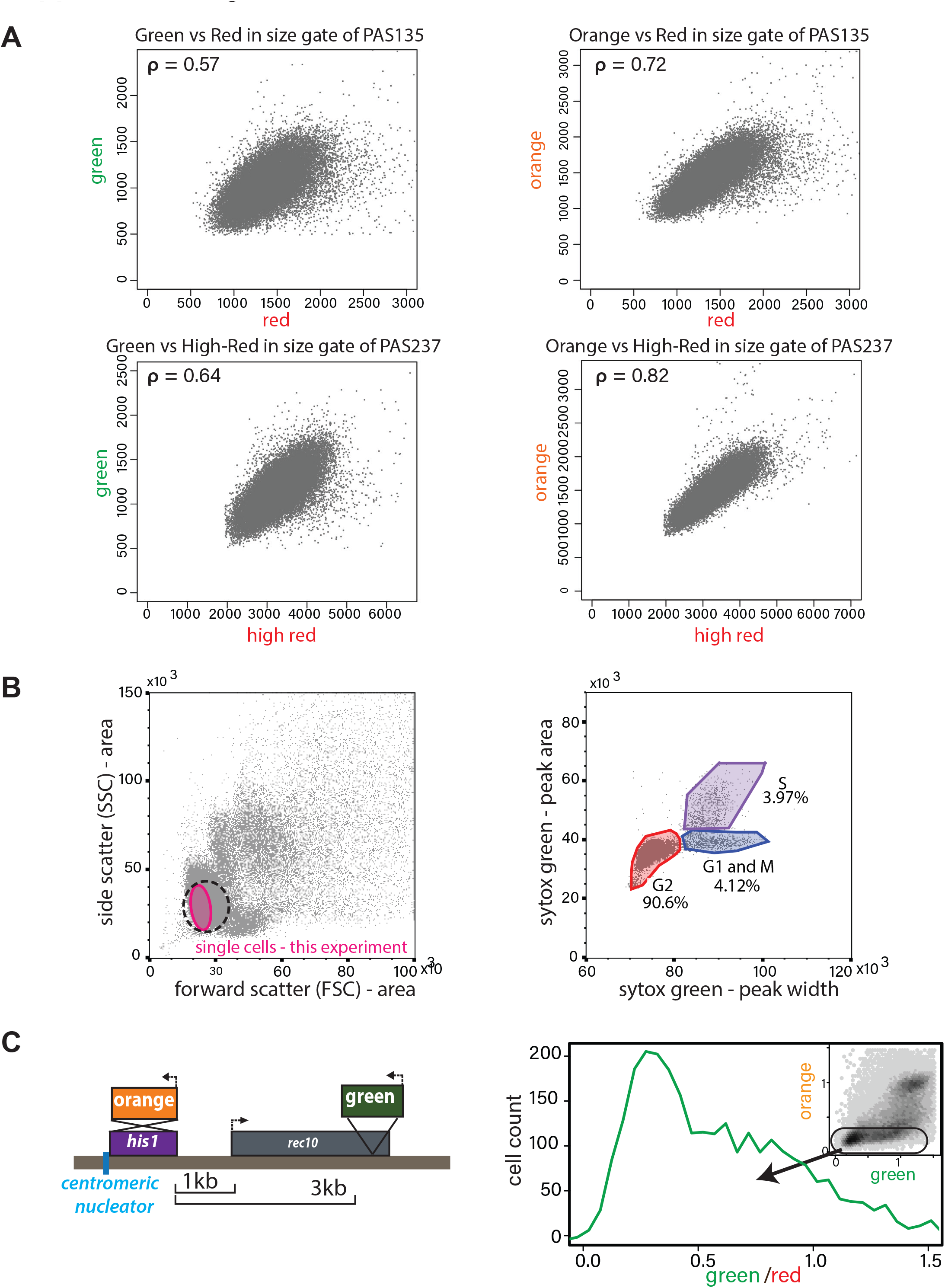
Validation of ectopic heterochromatin spreading sensor. **A.** Correlation of *ade6p:SFGFP* or *ade6p:mKO2* with *ade6p:3XE2C* (Red) or *act1p:1XE2C* (High Red) in *Δclr4* HSS size-gated (see B.) cells. LEFT: Plots of green vs. red channel signals of size-gated PAS 135 and 237 (both *Δclr4*, Red and High Red respectively). The Pearson correlation between “green” and “red” is shown. RIGHT: Plots of orange vs. red channel signals of size-gated strains as in LEFT. The Pearson correlation between “orange” and “red” is shown. **B.** Cell cycle stage of HSS and wild type cells by flow cytometry. Wild-type cells (PM03, see strain table) were fixed, stained with Sytox green DNA stain, and analyzed by flow cytometry. LEFT: side vs. forward scatter plot. Dotted line: The approximate size gate encompassing all experiments reported. Pink area: cells analyzed in the experiment shown. RIGHT: Plot of area vs. width parameter for the Sytox green channel, gates are drawn to denote cell cycle phases, G2 (red), G1 and M (Blue), S (purple) as described (Knutsen et al., 2011). **C.** Stochastic spreading and intermediate states produced by RNAi-driven nucleators are replicated at a second ectopic site. LEFT: Overview of the *his1::dh*HSS^3kb^. The colors are reversed relative to the *ura4::dh*HSS^1-7kb^ with “orange” as the “nucleation clamp” and “green” as the “sensor”. “Orange” replaces the *his1* gene and “green” is located 3kb downstream within the *rec10* open reading frame. RIGHT: histogram of “red”-normalized “green” fluorescence distribution of “orange”^OFF^ cells. Inset: 2D density hexbin plot.

**Supplemental Figure 2:**
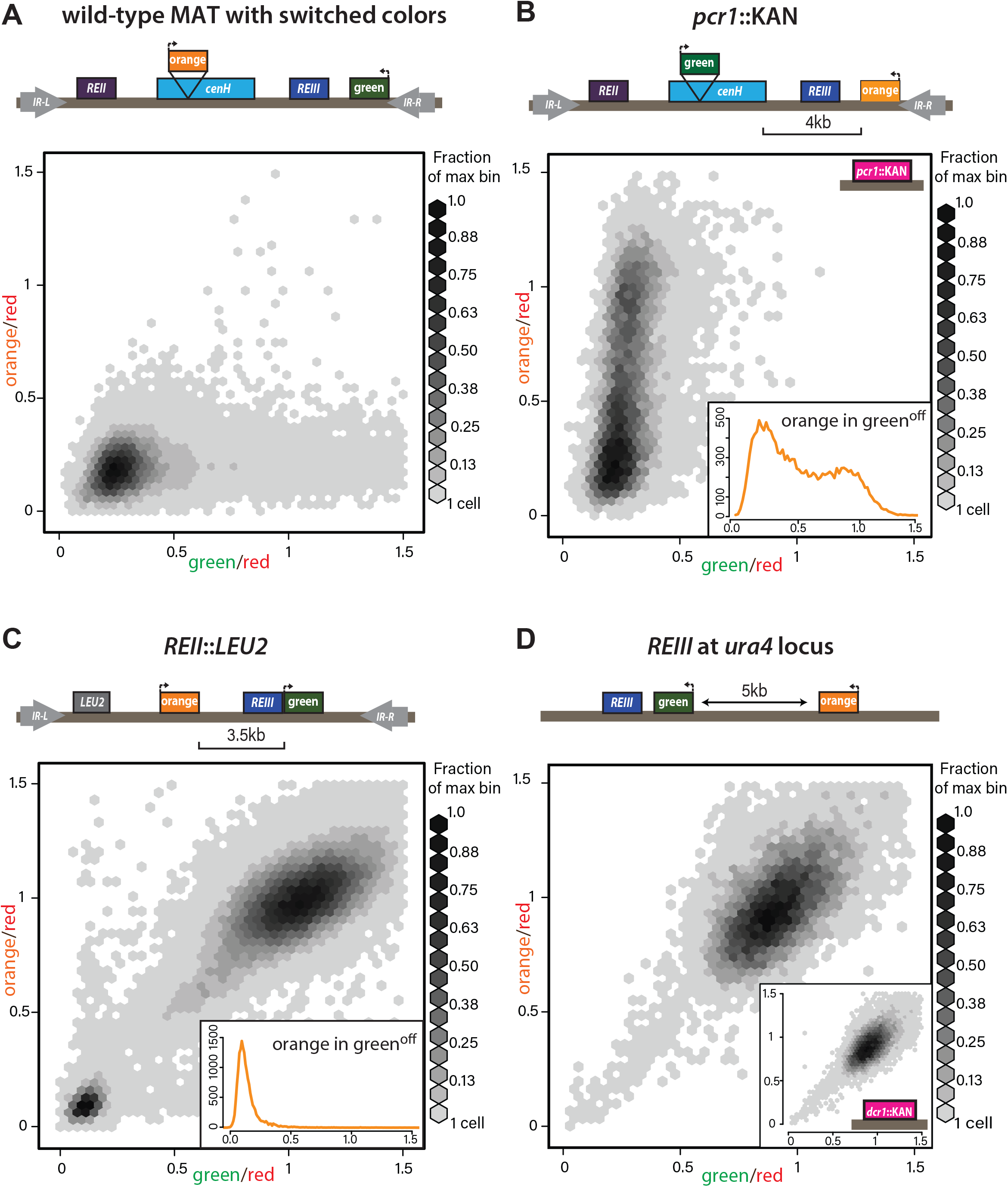
Heterochromatin spreading characteristics of nucleation elements at the tightly repressed MAT locus. **A.** The MAT^HSS^ documents tight repression of the wild type MAT locus. As in Figure 2A, with “green” and “orange” switched. The smallest bin starts with one cell and the maximal bin contains 429-458 cells. **B.** *pcr1::KAN* leads to stochastic spreading with intermediate states. *Pcr1* transcription factor was knocked-out in the PAS217 wild-type MAT^HSS^. Plot and inset as in Figure 2B, where the maximal bin contains 167-178 cells. **C.** *REII* does not contribute to bimodal distribution seen for *ΔK*^HSS^. The *REII* locus (1kb) was replaced with the *LEU2* gene before *clr4+* was introduced by cross. The smallest bin starts with one cell and the maximal bin contains 969-1034 cells. **D:** *REIII* is a weak nucleating element and unable to establish spreading at an ectopic site. 2D density hexbin plots of *ura4::REIIIHSS^5kb^* (PAS348), where the maximal bin contains 96-102 cells. Normalized green and orange are near 1.0, indicating a failure to repress both reporters. Inset: 2D density hexbin plots of *ura4::REIII*HSS^5kb^ *dcr1::KAN* (PAS350). *Dcr1* was deleted to release extra heterochromatin factors from RNAi-repressed loci. No additional silencing is detected.

**Supplemental Figure 3:**
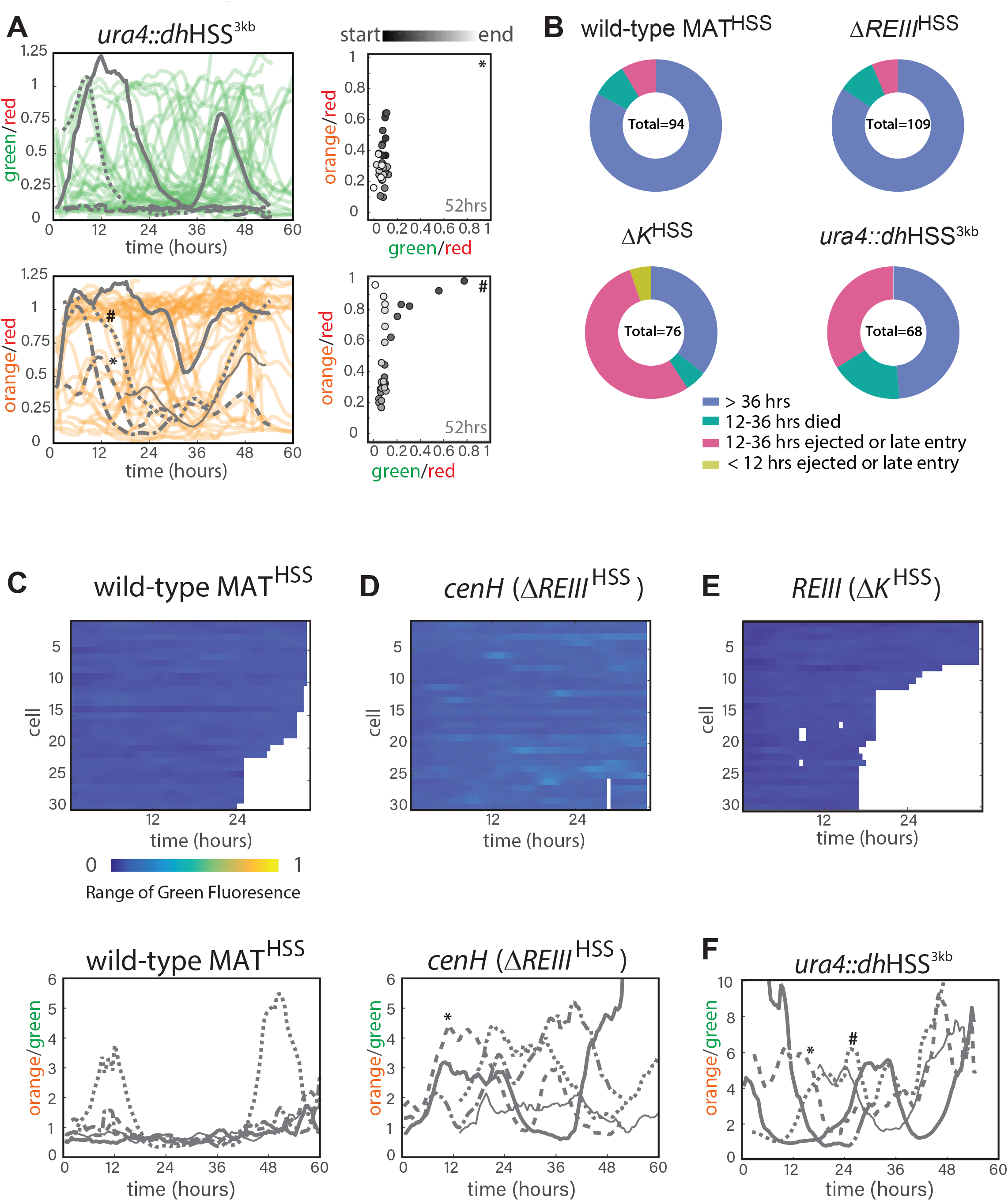
Single cell analysis of nucleation and spreading using a Fission Yeast Lifetime Machine (FYLM). **A.** FYLM analysis of *ura4::dh*HSS^3kb^ cells. TOP LEFT: 60hrs of normalized “green” fluorescence for all cells; 5 example cells are overlayed in gray each with different line types. BOTTOM LEFT: 60hrs of normalized “orange” fluorescence for all cells with the same 5 cells overlayed in gray. *, # represent two example cells. RIGHT: for two representative sample cells imaged, plots of normalized “green” and “orange” across its measured lifetime (grayscale). The corresponding cells are marked in the orange traces on LEFT. **B.** Categorization of cell longevity of all cells analyzed in the FLYM experiment. Measured lifespan ends when a cell dies or is ejected from its capture channel. **C.** For wild-type MAT^HSS^ TOP: “green” fluorescence heatmap (blue (0) to yellow (1)) for the same 30 cells as in 3C. BOTTOM: 60 hours of traces for “orange” divided by “green” for the five example cells indicated in 3C. **D.** “green” fluorescence heatmap and “orange”/”green” traces for *ΔREIII*^HSS^ as in C. **E.** “green” fluorescence heatmap *ΔK*^HSS^ as in C. **F.** “orange”/”green” traces for *ura4::dh*HSS^3kb^ as in C.

**Supplemental Figure 4:**
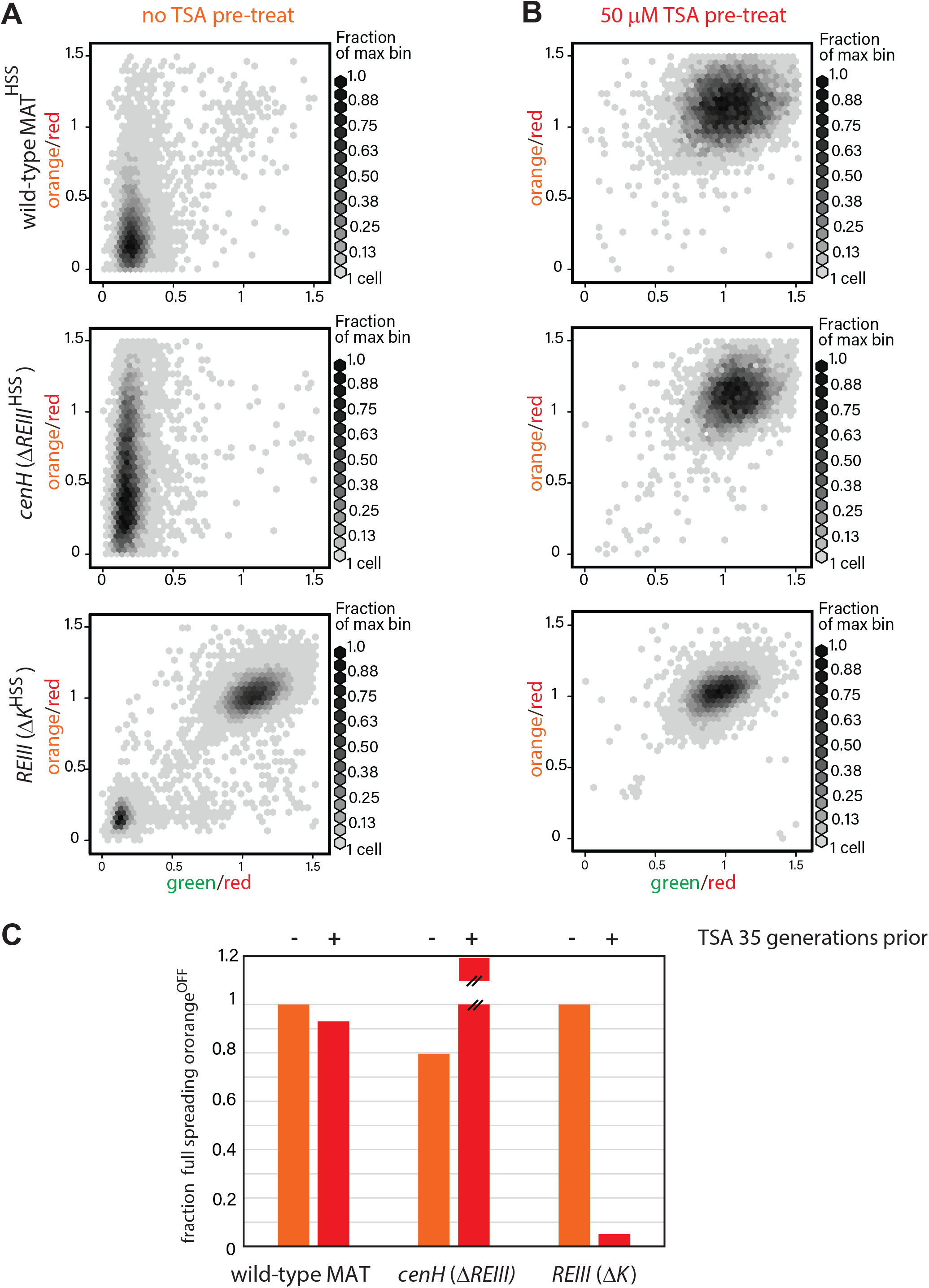
Behaviors of spreading during TSA treatment and after 35 generations. **A.** 2D density hexbin plots of wild-type MAT^HSS^, *ΔREIII*^HSS^, and *ΔK*^HSS^ strains grown 10 generations without TSA. For all panels, the smallest bin starts with one cell and the maximal bins from top to bottom contain 181-193, 97-103, 444-473 cells respectively. **B.** 2D density hexbin plots of wild-type MAT locus^HSS^, *ΔREIII*^HSS^, and *ΔK*^HSS^ strains grown 10 generations in 50 μM TSA. For all panels, the smallest bin starts with one cell and the maximal bins from top to bottom contain 63-67, 60-64, 197-210 cells respectively. The density distributions are near 1.0 in all strains indicating complete erasure of heterochromatin. **C.** History dependence at 35 generations after pretreatments. The fraction of cells with full spreading (wild-type MAT and *ΔREIII*) or fraction of cells with orange^OFF^ (*ΔK*) normalized to the highest value for ancestrally untreated cells (=1) is shown for the 0 μM TSA point. TSA pretreated cells for *ΔREIII*^HSS^ show higher repression than untreated cells. We interpret this to indicate experimental variations in silencing in the absence of memory. This is because for all other circumstances, TSA treatment results in reduced spreading, including for *ΔREIII*^HSS^ at 25 generations post treatment.

**Supplemental Figure 5:**
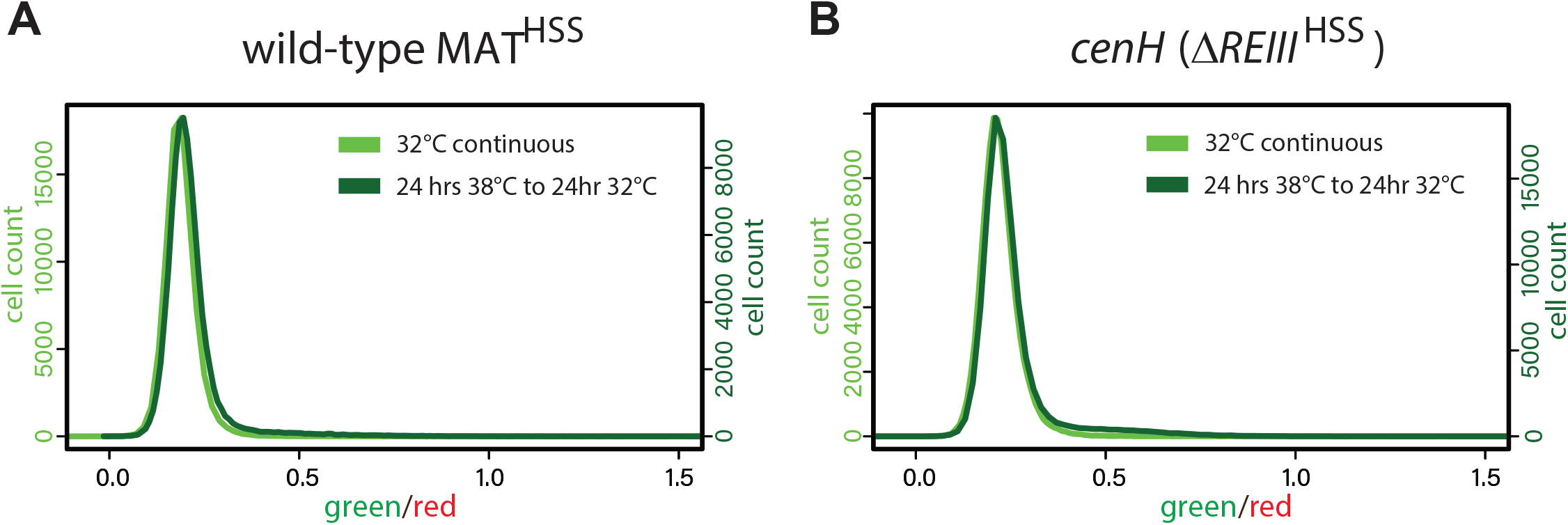
*cenH* nucleation is recovered within 24 hours at 32°C. **A.** 1-D histogram showing the distribution of green fluorescence in wild-type MAT locus^HSS^ cells grown either for 48hrs continuously at 32°C (left y-axis, light green) or heat stressed for 24hrs at 38°C followed by 24hrs growth at 32°C (right y-axis, dark green). **B.** For *ΔREIII*^HSS^ “green” histograms plotted as in A.

## Supplementary videos

**SVideo 1: Cell #274 from strain PAS244**

This movie consists of imaging in 4 channels, listed from top to bottom: Bright field, “green”, “orange”, and “red” for cell #274 from the strain PAS244 ura4HSS^3kb^. X-Y fluorescence plot for this cell is shown in Figure S3A, TOPRIGHT.

**SVideo 2: Cell #271 from strain PAS244**

This movie consists of imaging in 4 channels, listed from top to bottom: Bright field, “green”, “orange”, and “red” for cell #271 from the strain PAS244 ura4HSS^3kb^. X-Y fluorescence plot for this cell is shown in Figure S3A, BOTTOMRIGHT.

**SVideo 3: Cell #350 from strain PAS389**

This movie consists of imaging in 4 channels, listed from top to bottom: Bright field, “green”, “orange”, and “red” for cell #350 from the strain PAS389 WT MAT^HSS^. X-Y fluorescence plot for this cell is shown in Figure 3C.

**SVideo 4: Cell #407 from strain PAS391**

This movie consists of imaging in 4 channels, listed from top to bottom: Bright field, “green”, “orange”, and “red” for cell #407 from the strain PAS391 *ΔREIII*^HSS^. X-Y fluorescence plot for this cell is shown in Figure 3D.

**SVideo 5: Cell #123 from strain PAS387**

This movie consists of imaging in 4 channels, listed from top to bottom: Bright field, “green”, “orange”, and “red” for cell #123 from the strain PAS387 *δK*^HSS^. X-Y fluorescence plot for this cell is shown in Figure 3E.

## Supplemental Information

**1. Supplemental discussion**

**2. Supplemental references**

**3. Supplemental methods**

## Supplemental discussion

### Evidence in support of memory resulting from a TSA imprint at *REIII*

In principle, the difference between ancestrally TSA exposed and non-exposed wild-type MAT cells along the TSA concentration gradient (**Figure 4B**) could be explained by independent action of *cenH* and *REIII*. In this scenario, *REIII* does not re-nucleate following TSA pretreatment in wild-type cells, as is the case for *REIII* only (**Figure 4D**). Two lines of evidence argue against this model: First, the distribution of nucleated cells does not appear bimodal with respect to spreading, which would be expected in a population of *REIII* nucleated and non-nucleated cells (data not shown). Second, wild-type cells descended from TSA-treated ancestors are much more persistent in the TSA gradient than *ΔREIII* cells (**Figure 4B versus 4C**). Therefore, history dependence is unlikely to be accounted for simply by poor *REIII* re-nucleation. Further, we show that the presence of *cenH* dramatically increases *REIII* nucleation (see main discussion). Instead, we believe that TSA treatment leaves an imprint at the *REIII* locus that reduces its ability to stabilize spreading.

### Models for mechanistic differences in RNAi- and *REIII-* spreading and its inheritance

Our results show that spreading fidelity is dramatically increased for *REIII-* versus RNAi-mediated nucleation. Additionally, our results imply that when *REIII* heterochromatin is ancestrally erased, it is altered by a stable “imprint” that reduces *REIII*’s ability to support spreading in wild-type MAT (see above and main discussion). We propose two possible, but non-exclusive models to account for the molecular underpinnings of these observations: <u>Different heterochromatin composition.</u> *REIII*-induced spreading may contain different protein factors, which may stabilize the heterochromatin structure and modulate the efficiency of spreading. It is already understood that certain factors are recruited differentially and/or loaded by multiple mechanisms at the different endogenous heterochromatin elements in the cell (Cam et al., 2005; Ekwall et al., 1999; Petrie et al., 2005; Sugiyama et al., 2007). Some of those may act for example to modulate the production of the H3K9me3 mark by Suv39/Clr4, which is critical for spreading (Al-Sady et al., 2013; Jih et al., 2017). In support of this model, the loss of Clr3, which is directly recruited by Atf1/Pcr1 to *REIII*, reduces H3K9me3 (Yamada et al., 2005). Persistence in the face of perturbations as well as high likelihood of inheriting positional information in the next generation, could be achieved if this specialized heterochromatin is protected from histone loss and demethylation by Epe1. *REIII* and *cenH* spreading may attract different concentrations of factors described to antagonize histone loss (Taneja et al., 2017; Yamane et al., 2011) and/or antagonize Epe1 (Braun et al., 2011). An imprint at *REIII* by TSA treatment could be explained by opposing antagonistic positive feedback loops, which are commonly associated with hysteresis, or memory, in cellular reactions (Bagowski and Ferrell, 2001). One possible example of a molecular implementation is maintenance of reduced nucleosome occupancy: The loss of components of the SHREC complex, some of which are directly recruited at *REIII* by Atf1/Pcr1 (Kim et al., 2004), induce nucleosome free regions (NFRs) near *REIII* (Garcia et al., 2010). NFRs or low nucleosome occupancy in general is known to disfavor spreading (Garcia et al., 2010; Yamane et al., 2011). If, next to *REIII*, a TSA-induced NFR or lower nucleosome occupancy is maintained by positive feedback, it could serve as a memory imprint. <u>Specialized three dimensional structure.</u> It is conceivable that *REIII*-but not RNAi-mediated heterochromatin spreading adopts a specialized and highly stable structure, distinct from models for the isolation of overall heterochromatin sites, such as the entire MAT locus (Noma et al., 2001). The formation of a local structure is consistent with our findings and those of others (Wang and Moazed, 2017) that *REIII-* nucleated spreading, unlike RNAi-spreading (**Figure 1C, 2B** and **S1C**), cannot be re-capitulated outside MAT at the *ura4* locus (**Figure S2D**), even when additional heterochromatin factors are made available by compromising RNAi-nucleation (**Figure S2D, inset**). If this structure were for example a compact loop, only across a few kilobases, it would account for deterministic spreading, as looping would facilitate reaching all nucleosomes within this structure, as has been proposed (Bantignies and Cavalli, 2011). In this model, memory would not require retention of methylated histones, as there would be a strong bias to reconstitute spreading after S-phase at the locus if the structure was formed. But memory would require a strong bias to replicate this structure after S-phase with history dependence also resulting from a significant energetic barrier to form the structure *de novo*. Given this postulated barrier to loop formation, a TSA imprint would simply require TSA to favor dissolution of the loop, which would then be maintained and weaken spreading. Mechanisms of loop formation are poorly understood in *S. pombe*, but may involve the chromatin organizing TFIIIC complex, which is recruited as part of the spreading boundary adjacent to *REIII* (Noma et al., 2006). In support of this overall model, looping in heterochromatin spreading has been predicted to lead to the more robust inter-generational persistence that we witness here (Erdel and Greene, 2016) for REIII-mediated spreading.

### Alternative formal model for *cenH* and *REIII* interaction

In the main discussion, we propose that *cenH* stimulates *REIII* nucleation to account for the high proportion of the spreading in wild-type MAT cells. Another formal possibility remains that non-nucleated, Atf1/Pcr1-bound, *REIII* raises *cenH* spreading efficiency. In the *ΔREIII* strain, Atf1/Pcr1 binding sites have been fully deleted, it is possible that when bound but not in a heterochromatin state, Atf1/Pcr1 act in some manner to encourage more efficient spreading out of *cenH*, possibly by directing the locus to a more spreading competent location.

## Supplemental Methods

### Basic 3 color HSS Analysis in R

#### Reading in the data

Standard flow cytometry data files (.fcs) were converted to readable “dataframes” (spreadsheets) using the read.FCS() and exprs() functions in the R package flowCORE (Bioconductor, https://www.bioconductor.org/packages/release/bioc/html/flowCore.html).

#### Generating cell size gate

Forward Scatter Area (FSC) and Side Scatter Area (SSC) parameters were extracted for each sample and a scatter plot of FSC vs SSC was generated. As shown in **Figure S1B**, a gate was determined for small single cells of predominantly G2 phase. A function was created to isolate the data in the dataframe for each sample within the FSC and SSC parameters identified.

#### Isolating successfully nucleated cells

For all flow cytometry experiments done at standard temperature and without TSA, successfully nucleated cells were determined as follows. A strain closely matched in genetic background to HSS strains but containing no XFPs was analyzed under the same flow cytometry conditions in each experiment. This “unstained” control was gated for cell size in the same manner as analysis strains and both the mean fluorescence and standard deviation determined in Green or Orange Channels (the signal from the *ade6p:*SF-GFP, *ade6p:*mKO2 or *ade6p:*E2C transcriptional units is referred to here as “green”, “orange” or “red”). A nucleation cutoff (“green” for all experiments except **S1C** and **S2A**) was set for a value corresponding to the mean of fluorescence units in the green channel plus two standard deviations from this unstained control. Only cells for each analysis strain having a “green” (or “orange” in **S1C, S2A**) signal less than this value were considered for post nucleation analysis.

#### Normalizing to max fluorescence values from Δclr4 strains

Max values in *Δclr4* strains were determined by calculating the median raw fluorescence in each color channel after gating for cell size. For each cell of each strain for analysis, the signal in each channel was divided by this max value for the corresponding *Δclr4* strain.

#### Normalizing to constitutive red signal

After normalizing to the max values in each channel, for each cell of each strain, the “green” and “orange” values were divided by the “red” value. The normalized values range from 0 to 1, where 1 corresponds to the *Δclr4* (max) value. As this value is derived from the mean of a cell distribution, with *Δclr4* cells falling above and below the mean, we plotted out to 1.5 to capture cells with ratio values above 1.0.

#### Hexbin (2-D Histogram) Analysis

All cells within the FSC/SSC gate were plotted as a “green”/“red” and “orange”/“red” hexbin (or 2-D histogram) plot where density is color-coded in grayscale. Data points within x=0-1.5 and y=0-1.5 were isolated and a hexbin plot was generated using n=40 bins. Hexbin plots were generated using the R package hexbin (https://CRAN.R-project.org/package=hexbin)

#### Spreading Analysis with Nucleation Clamp

Cells within the FSC/SSC gate and with a nucleation color signal below the cutoff value were plotted in a 1-D histogram where the points in the middle of the histogram bins were plotted connected by a line. In **Figure 1**, 1-D histograms were generated using n=100 bins. In the remaining figures where more data was collected or where there were more cells within the nucleation gate plots were generated with n=200-300 bins.

### Modifications for S1 A

After reading in the .fcs file and gating for cell size, the raw fluorescence in Green vs Red and Orange vs Red were plotted for each cell in either PAS135 or PAS237. Color negative cells were removed. Pearson correlation for each color pair in each strain was calculated.

### Modifications for Figure 4

For every TSA concentration and pre-condition (0 μM or 50 μM) each strain was normalized to the median fluorescence of a *Δclr4* strain grown under the same treatment. Both “green” and “orange” cutoff values for each analysis strain were generated by determining mean and two standard deviations in each color from PAS217 at 0 μM TSA from the 0 μM TSA precondition normalized to the appropriate *Δclr4* strain for each analysis strain. Previous analysis had demonstrated both colors in PAS217 to be fully repressed, as evident in the mean for each channel. Values for “green”/“red” and “orange”/“red” were divided by “green”/“red” and “orange”/“red” values for each *Δclr4* strain at each TSA concentration and pre-condition, yielding the cutoff value as described above. For **4B** and **4C** we calculated at each TSA concentration the fraction of cells with green signal below the “green” cutoff that have an orange signal below the “orange” cutoff. This yields a fraction of nucleated cells which have “full spreading” as measured by the sensitivity of our reporters. For **4D** we calculated at each concentration of TSA the fraction of all cells with orange signal below the “orange” cutoff, as in the 50 μM TSA pre-condition, insufficient cells exist that are below the cutoff for “green” to perform above analysis. This line plotted however is co-incident with the fraction of all cells with “green” signal below the “green” cutoff along the TSA concentration gradient (data not shown).

### Modifications for Figure 5

In **Figure 5A** for each temperature a different FSC/SSC gate was determined as the size and shape of cells are affected by changes in temperature. A corresponding “red only” (PAS75) was also grown at that temperature and subjected to the same FSC/SSC gate. Nucleation (green) and spreading (orange) cutoff values were determined based on the ratios of this “red only” strain normalized to *Δclr4*. For each strain at each temperature, we calculated the fraction of all cells in the FSC/SSC gate that had “green” signal less than the “green” cutoff AND “orange” signal less than the “orange” cutoff. These values were normalized to the fraction calculated for cells of that strain at 32°C.

In **Figure 5 C, D, E** the 1-D histograms and hexbin plot insets were calculated as above with the modification that the “green” cutoff values were generated as in **Figure 5A**.

In **Figure 5F** we calculated the fraction of gated cells that had “green” signal less than the “green” cutoff AND orange signal less than the “orange” cutoff. This value was normalized a similar fraction calculated for that strain at 32°C on day 0 of the experiment.

### Fitting t_1/2_ for 38°C spreading recovery

To derive a t_1/2_, which is the time required to recover to 50% of the full spreading observed at 32°C, we fit the data to a simple sigmoidal dose-response variable slope model:

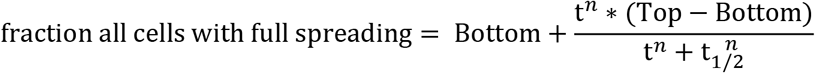

where Bottom is the starting fraction of cells with full spreading at t=0, Top =1, t is time in hrs. *n* represents a Hillslope.

### FYLM data analysis

#### Initial data calculations

Loss of focus was identified by red fluorescence measurements below a cutoff of one standard deviation from the mean of all collected values of red for all cells. This loss of focus data was removed from analysis. Background fluorescence from the PDMS device at each time point was then subtracted using catch tubes that did not receive a cell. For each MAT strain, its matched *Δclr4* strain was also imaged and a mean and standard deviation were calculated. In each strain cells were normalized to this mean *Δclr4* value (defined as “max”) and to their own red values as in the flow cytometry data analysis. An ON gate (used in **Figure 3B**) for cells that reached maximal de-repression was calculated for each strain from the *Δclr4* strain mean less 3 standard deviations. For the ectopic locus strain where both colors reach full derepression at various points throughout the experiment, the “max” was calculated differently. For each cell of this strain green and orange values were each divided by the red value at every timepoint. To determine max = 1 for orange, a Gaussian mixture model with two components was fit to an array of all of the orange/red values. The higher mean of was used as the “max” value and all orange data points were normalized to this. For green “max” was set to the maximum green/red value measured within the first 12 hours of the experiment.

#### Calculating nucleation gates

As seen by flow cytometry and visual inspection of collected movies, the vast majority of MAT locus cells have a repressed nucleation reporter (green), which allowed us to formulate a very strict nucleation cutoff from the collected FYLM data itself. This cutoff was the mean plus two standard deviations of all measured values of all cells. Only cells that maintained a green signal less than this cutoff for their entire measured lifespan were included for further analysis in Figure 3. We did not apply this nucleation gate to the ectopic strain, as only 2 cells maintained “green” tightly repressed throughout their measured lifespan. Instead, we decided to show all the cells in the traces plot and highlight in grey example cells that remain nucleated or mostly nucleated throughout their measure lifespans.

#### Rescaling orange to fix negative values

Due to background subtraction (see above) a significant fraction of cells experienced negative values for their adjusted fluorescence in the orange channel. To account for this, the data was rescaled by determining the lowest value measured (minVal) and adding the difference between that value and 0 to every timepoint of every cell. Values for all timepoints were then divided by 1+minVal to rescale back to 1 = max.

#### Data smoothing

For trace plots and heatmaps data was smoothed using a moving average of the two-nearest neighbor data points before and after. This number was chosen as it represents the timeframe of one cell division and is on the timescale of the expression and maturation of the XFPs used in these strains.

#### Traces

Individual cell traces represent the red normalized and smoothed, green and orange fluorescence data plotted over time. Traces begin and end at whatever time a cell entered or exited the channel or died. Therefore, not all traces start at x = 0 or end at x = 60. Curated example cells were also plotted as overlays using gray lines. For these curated cells similar trace plots for orange divided by green was plotted in **Figure S3C BOTTOM, S3D BOTTOM and S3F**.

#### Heatmaps

Points with red values greater than 50% of the mean were removed. For cells that remained nucleated throughout their measured lifespan, up to 36 hours of measurements of normalized green or orange was plotted as a heat map from blue (0) to yellow (1) for 30 of the longest imaged cells. White gaps indicate transient loss of focus of less than 2 hours (4 timepoints). Heatmaps were no longer plotted for any cell that had a loss of focus event for more than 4 timepoints.

#### X-Y fluorescence plots

For one or two selected cells per strain an X-Y fluorescence plot was generated that plots the orange vs green values for every third timepoint imaged. Points are colored in a grayscale that is generated based on the measured lifespan of that cell. The first measured point is represented in black, the last in white and the number of remaining points set by the total measured lifespan of the cell.

#### Cell fate pie charts

The number of measured timepoints for each cell was determined and converted to hours (1 fluorescence image per 30 minutes). Cells were then binned into lifetime groups of <12hrs, 12-36hrs, or >36hours. Within these bins the cells were separated based on whether they died as annotated in FYLM Critic or if their traces were cut short due to late entry into or ejection from the catch channel.

